# Multiomic Profiling Links RDH12-Dependent Retinaldehyde Detoxification to Membrane Remodeling and Ferroptosis

**DOI:** 10.64898/2026.09.24.754135

**Authors:** KanagaVijayan Dhanabalan, Miranda R. Muhoberac, Ramya Visvanathan, Devender Arora, Olga Belyaeva, Natalia Y. Kedishvili, Ramaswamy Subramanian, Bryon S. Drown

## Abstract

All-*trans*-retinal (atRAL) is a photoreactive aldehyde generated during visual pigment regeneration, and impaired atRAL clearance contributes to oxidative stress and retinal degeneration. Retinol dehydrogenase 12 (RDH12) reduces atRAL to all-trans-retinol, yet the temporal mechanisms linking its upstream enzymatic activity to downstream cellular stress adaptations remain poorly defined. Here, we utilized stable RDH12-expressing HEK293T cells to match proteomic and lipidomic profiles and performed cell viability assays to map the transition from acute atRAL stress to cellular recovery. RDH12 expression increased cellular tolerance to retinaldehyde-induced cytotoxicity. Acute atRAL exposure engaged the supply arm of the KEAP1-NRF2 program with increased HMOX1, SLC7A11, GCLC, MGST2, and MGST3, while the terminal glutathione peroxidase arm declined. Acute exposure was accompanied by membrane remodeling, indicated by increased glycerophospholipid enzyme abundance and increased stress-signaling ceramides and lysophosphatidylcholines. Concurrently, acute stress induced upregulation of the transferrin receptor (TFRC) and SLC11A2, which influence intracellular iron flux and enable Fenton chemistry that can lead to oxidative stress. During recovery, RDH12-expressing cells exhibited a distinct post-atRAL molecular state characterized by increased abundance of phosphatidylcholines (PCs) and ether-linked PCs, and proteome changes associated with cytoskeletal organization and antioxidant processes. Furthermore, we performed a pharmacological rescue with Ferrostatin-1, which indicated that atRAL-induced cytotoxicity in RDH12-expressing cells is dependent, in part, on lipid radical propagation and lipid peroxidation, supporting ferroptosis-regulated cell death. Collectively, the multiomics and biological findings establish a novel temporal sequence for retinaldehyde toxicity, showing that RDH12 is a critical component of upstream metabolic defense for acute aldehyde clearance, late-stage lipid detoxification, and adaptive membrane lipid homeostasis.

## Introduction

Photoreceptor degeneration is a hallmark of retinal degenerative diseases (RDDs), a group of disorders driven by chronic oxidative stress (1). Photoreceptors are highly vulnerable to oxidative damage owing to high concentrations of unsaturated lipids, high metabolic activity, and continuous recycling of all-*trans*-retinal (atRAL), a highly reactive and cytotoxic aldehyde essential for vision (2). Following photoactivation of rhodopsin, 11-cis-retinal is converted to atRAL and subsequently recycled to all-*trans*-retinol as part of the visual cycle (2, 3). Impaired atRAL clearance contributes to oxidative stress through reactive oxygen species production, lipid peroxidation, and accumulation of misfolded proteins, ultimately triggering cell death (4–7). Thus, efficient atRAL clearance is fundamental to photoreceptor longevity (8). Defective all-*trans*-retinal clearance has recently been shown to drive photoreceptor degeneration through ferroptosis in mice (9), providing a direct link between retinaldehyde overload and lipid-peroxidation-dependent cell death.

Excessive or prolonged atRAL accumulation can activate oxidative stress and cell-death processes. Because atRAL contains a highly reactive aldehyde group, it readily forms Schiff-base and Michael adducts with protein and phospholipid amines, seeding bisretinoid formation and protein modification (10, 11). In parallel, atRAL promotes oxidative stress through reactive oxygen species generation and lipid peroxidation (12). The lipid composition of photoreceptor membranes is enriched in docosahexaenoic acid DHA and very long-chain polyunsaturated fatty acids VLC-PUFAs (13, 14). This high degree of unsaturation is essential for the fluidity, thickness, and functional properties of outer-segment membranes; however, these lipids are susceptible to oxidation by free radicals (15, 16). Additionally, atRAL exposure has been associated with endoplasmic reticulum stress, mitochondrial dysfunction, stress-kinase signaling, and oxidative damage (6, 17). Recent studies have implicated ferroptosis, an iron-dependent form of regulated cell death driven by lipid peroxidation in photoreceptors (9). Inhibition of JNK signaling has been shown to attenuate atRAL-induced photoreceptor ferroptosis, linking atRAL upstream of stress-kinase and lipid-peroxidation-dependent degeneration (18). AtRAL has also been linked to JNK-mediated apoptosis and Gasdermin E (GSDME)-mediated damage, and GSDME activation has been shown to increase photoreceptor sensitivity to ferroptosis (17, 19). Together, these studies suggest that retinaldehyde accumulation lies upstream of interconnected electrophilic-stress, membrane-damage, and regulated cell-death pathways.

Retinaldehyde metabolism is carried out by NADPH-dependent retinol dehydrogenases (RDHs) that catalyze the oxidation and reduction reactions required for visual cycle function within photoreceptors and the retinal pigment epithelium. Among these enzymes, RDH8, RDH11, and RDH12 are involved in the reduction of atRAL to all-trans-retinol, thereby limiting the accumulation of this highly reactive aldehyde following photoactivation. Mutations in these enzymes have been genetically linked to severe RDDs, including cone-rod dystrophy 2, Leber’s congenital amaurosis 13, retinitis pigmentosa, and age-related macular degeneration. RDH12 is an NADPH-dependent reductase that converts atRAL to all-trans-retinol (atROL) within the inner segments of photoreceptor cells (20–22). Pathogenic variants in the RDH12 gene cause early-onset, severe retinal dystrophy characterized by rapid, progressive photoreceptor loss (21, 23, 24). The visual function in patients with mutations in RDH12 or RPE65, another LCA gene involved in retinoid metabolism, shows distinctly different pathologies (25–28). Mouse models lacking RDH12 show delayed dark adaptation, accumulation of atRAL after light exposure, and increased susceptibility to light-induced retinal degeneration (20, 29). RDH8 and RDH12 double knockout substantially diminishes all-trans RDH activity, with RDH12 providing protection against retinaldehyde. Despite its clinical relevance, the cellular mechanisms by which RDH12 modulates the stress response against reactive aldehydes remain poorly understood. RDH12 has traditionally been viewed as a metabolic enzyme that limits aldehyde accumulation; however, its broader role in orchestrating cellular stress responses, organelle homeostasis, and lipidome remodeling remains unclear.

Prior transcriptomic and targeted metabolomic studies have provided valuable insights into retinal degeneration (30–33). Previous systems-level investigations have primarily utilized *Abca4−/−* and *Abca4−/− Rdh8−/−* mouse models, alongside ARPE-19 or 661W cell lines, focusing predominantly on the chronic accumulation of bisretinoids (such as A2E) and late-stage disease pathology (4, 8, 34–37). These studies have successfully highlighted broad markers of oxidative stress, mitochondrial dysfunction, and generalized lipid peroxidation (4, 35, 37, 38). Most prior analyses capture static snapshots of chronic damage rather than the dynamic, temporal shifts that occur during acute atRAL stress and subsequent cellular recovery. Furthermore, while RDH12 is genetically recognized as a critical defense enzyme, the global molecular network by which it orchestrates downstream stress adaptations, specifically mapping its protective influence across the proteome and lipidome, has not been systematically characterized (5, 20, 26, 39, 40).

In this study, we hypothesized that RDH12 expression fundamentally alters the cellular threshold for atRAL-induced toxicity and changes the molecular state from acute stress to recovery. To capture this temporal change, we employed multi-omics, proteome and lipidome profiling in a stable RDH12-expressing HEK293T cell line across two experimental conditions: an acute atRAL exposure phase and a subsequent recovery phase (**Figure 1A**). Our analyses enabled us to examine dose-dependent cytotoxicity and changes in proteostasis, membrane lipid metabolism, the lipidome, and ferroptosis-regulated cell death (**Figure 1B**). RDH12 expression increased cellular tolerance to atRAL and was associated with a distinct proteome and lipidome state during recovery. Pharmacological rescue with Ferrostatin-1 supported lipid peroxidation-dependent cell death in response to atRAL-induced toxicity. Our results show RDH12 is an important enzyme in the response to retinaldehyde stress and the cellular processes that influence membrane recovery.

**Figure 1.**
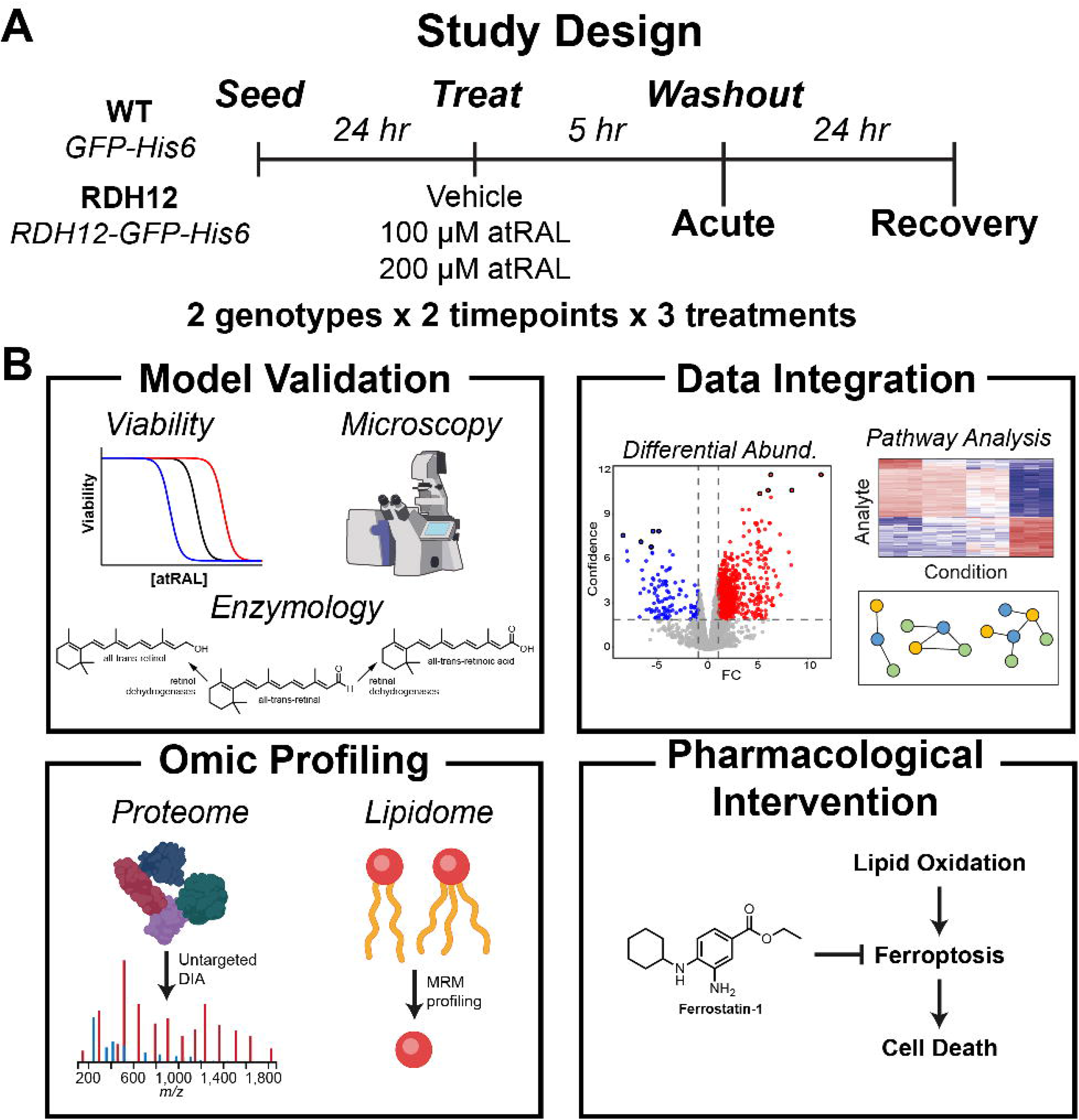
Study design and overview of measurement panel. *A*, HEK293T cells stably expressing either unconjugated GFP (WT) or RDH12-GFP fusion (RDH12). Cells were transiently treated with atRAL and collected after 5 h of treatment (acute) or 24 h after compound washout (recovery). Each treatment and time point was conducted with three independent biological replicates. *B*, Phenotypic validation of the cell model included dose-dependent viability, subcellular localization, and measurement of specific activity. Paired proteomic and lipidomic analyses captured concerted response to atRAL toxicity and recovery. After integrating the data, we pharmacologically evaluated the ferroptotic cell death mechanism.

### Experimental Procedures

#### Experimental Design and Statistical Rationale

Two label-free DIA proteomics experiments were performed on HEK293T cells. In the acute experiment, RDH12-expressing cells were treated for 5 h with vehicle (ethanol), 100 µM atRAL, or 200 µM atRAL (three conditions). In the recovery experiment, RDH12-expressing and WT (GFP) cells were treated for 5 h with vehicle, 100 µM, or 200 µM atRAL, washed, and allowed to recover for 24 h (2 genotypes × 3 treatments = six conditions). Every condition comprised three independent biological replicates, each a separately seeded, treated, and harvested T25 flask, giving 9 acute and 18 recovery samples (27 in total). Each sample was injected once (no technical replicates) on a Thermo Orbitrap Exploris 480, for 27 single-shot DIA runs. A separate basal comparison of untreated RDH12-expressing and WT cells (n = 3 biological replicates per genotype, six runs) was acquired on a Bruker timsTOF HT to assess the proteomic consequence of RDH12 overexpression (**Figure S1**). These data were not combined with the acute or recovery datasets. Recovery samples were injected in randomized order, and acute samples were injected in an interleaved order (vehicle and 100 µM atRAL alternating by replicate, followed by the 200 µM replicates). No retention-time standards or spiked protein or peptide standards were used. MS1 spectra were acquired in every cycle; the precursor m/z range was fractionated into 52 variable-width isolation windows (12 or 24 m/z, 0.5 Th overlap, SI Table 1) with a total cycle time of approximately 3.1 s. No sample-derived spectral library was generated from DDA as an in silico predicted library was used and refined in a first pass over the 27 DIA runs themselves (see Data processing and analysis).

Sample size was chosen to provide three biological replicates per condition, the minimum required for moderated t-statistics, and a fully crossed genotype x treatment design in the recovery experiment. Differential protein abundance was tested with the Bioconductor package DEP (41), which fits a linear model to each protein and applies limma empirical-Bayes moderated t-tests (42). Moderation borrows variance information across the ∼8,000 quantified proteins and is suited to the small number of replicates per condition. P-values within each contrast were adjusted by the Benjamini-Hochberg procedure to control the false discovery rate at the protein level. Proteins were called differentially abundant at |log_2_ fold change| ≥ 1 and adjusted p < 0.01 for the volcano plots and GO enrichment. The pathway heatmaps use a relaxed display threshold of |fold change| ≥ 1.4 with adjusted p < 0.1 to show the direction of coordinated pathway members.

Before testing, proteins were required to be quantified in at least two of three replicates in at least one condition, and remaining missing values were imputed with a mixed strategy (k-nearest neighbors for values classified as missing at random and QRILC for values missing not at random). Contrasts in which neither condition had an observed value were excluded from testing, as described under Data processing and analysis. Reproducibility was assessed from protein-level coefficients of variation across biological replicates (non-imputed intensities, ), identification overlap across replicates, and principal component analysis (**Figure S2**). Cell-viability dose– response experiments (n = 3 biological replicates) were analyzed as described under Cell Viability and Ferroptosis Inhibition Assays.

#### Generation of Stable RDH12-Expressing Cell Lines

The human RDH12 open reading frame (UniProt: Q96NR8) was cloned into the pLVX-Puro lentiviral expression vector (VectorBuilder) with a C-terminal GFP followed by a 6xHis-tag. Lentiviral particles were produced by VectorBuilder. We transduced target HEK293T cells with viral supernatants in the presence of 8 µg/mL polybrene. At 48 hours post-transduction, we selected stable transformants under puromycin pressure (3 µg/mL) for 3 days. Cells were maintained in the presence of puromycin thereafter. We verified RDH12 expression by immunoblotting and confocal microscopy (RDH12). We have generated HEK293T cells expressing GFP with a 6X His-tag as a control line using the same method described above (WT).

#### Cell Culture

HEK293T cells were maintained in Dulbecco’s Modified Eagle Medium (DMEM) supplemented with 10% Fetal Bovine Serum (FBS) and 1% Penicillin-Streptomycin. Cells were cultured at 37 °C in a humidified atmosphere containing 5% CO_2_. All-trans-retinal (atRAL; Sigma-Aldrich) was prepared as a 1 mM stock in absolute ethanol and then diluted in complete media. To prevent photo-oxidation and isomerization, we prepared all atRAL stocks and treatment media fresh under dim red light and handled them in amber tubes.

#### Cell Viability and Ferroptosis Inhibition Assays

Cell viability was evaluated using the CellTiter-Glo Luminescent Cell Viability Assay (Promega – G9241). Cells stably expressing RDH12-GFP or a GFP control were seeded into 96-well plates at a density of 5,000–10,000 cells per well. After 24 hours, cells were challenged with 0–400 µM atRAL for 5 hours. For ferroptosis inhibition experiments, cells were pretreated with 30 µM Ferrostatin-1 (Fer-1) for 24 hours before atRAL exposure. After treatment, we added CellTiter-Glo reagent according to the manufacturer’s protocol and recorded luminescence on a microplate reader. Dose-response curves were fitted with a four-parameter log-logistic model (drc::LL.4) with the upper and lower asymptotes fixed at 100% and 0% respectively. IC_50_ values are pooled fits across replicates with 95% confidence intervals. Per-replicate IC_50_ values were compared by two-factor ANOVA on log_10_(IC_50_) with Tukey HSD.

Sample Collection for Multiomic Analysis: RDH12-GFP and GFP cells were seeded into T25 flasks at a density of 0.8 × 10⁶ cells per flask. After 24 hours, cells were treated with 100 µM or 200 µM atRAL for 5 hours. Ethanol-treated cells served as a vehicle control. To evaluate the recovery phase, treated cells were washed with 1X PBS after the initial 5-hour atRAL exposure, replenished with fresh DMEM containing 10% FBS, and incubated for an additional 24 hours. At the designated endpoints, viable cells were washed with 1X PBS, harvested by scraping, snap-frozen in liquid nitrogen, and stored as pellets at -80 °C until analysis.

#### Confocal Microscopy

Cells stably expressing RDH12-GFP or GFP were seeded onto glass-bottom coverslips and grown to 60–70% confluency. Add ER-Tracker Red and incubate for 30 minutes. Cells were then fixed with 4% paraformaldehyde (PFA) for 15 minutes at room temperature. After three PBS washes, we mounted the coverslips onto slides. We acquired fluorescence images using a Nikon A1Rsi laser-scanning confocal microscope equipped with a 100× oil-immersion objective (NA 1.49). We imaged samples under identical acquisition settings whenever we compared experimental groups. We used sequential scanning to minimize spectral overlap between fluorescence channels. We performed image acquisition and instrument control using NIS-Elements software (Nikon). We collected images as needed and processed them using Nikon NIS-Elements software. Final images were adjusted uniformly for brightness and contrast across all samples without altering the original fluorescence distribution.

#### Enzyme Activity Assay

To evaluate RDH12 activity in intact cells, HEK293T cells stably expressing RDH12-GFP (RDH12) or a GFP control (WT) were plated in 6-well dishes. The next day, nearly confluent cells were treated with 5 µM all-*trans*-retinal (atRAL, Toronto Research Chemicals, #TRC-R240000) for 4 hours. atRAL stock was prepared in ethanol and added to the culture medium with a final ethanol concentration not exceeding 0.3%. Following the incubation, cells and culture medium were harvested separately. Total cellular protein content in each well was determined using DC Protein Assay II (BioRad Laboratories, #5000112). Retinoids from the culture medium were extracted in hexane and analyzed by normal-phase HPLC, essentially as described before (43) using Waters 2695 Separation module equipped with Waters 2998 PDA Detector. The mobile phase was hexane/ethyl acetate/acetic acid (95:4.975:0.025, v/v/v) at a flow rate of 0.7 ml/min. The stationary phase was Waters Spherisorb S3W column (4.6 × 100 mm). Retinoids were quantified by comparing their peak areas to a calibration curve constructed from peak areas of a series of standards. The amount of retinoids produced in each culture well was normalized by the total cellular protein content in the well.

#### Immunoblotting and Protein Analysis

Cells were lysed in RIPA buffer supplemented with protease inhibitor cocktails, and total protein concentrations were determined using a BCA assay. Equivalent amounts of protein (20–30 µg) were resolved on 12% SDS-PAGE gels and transferred to PVDF membranes. To detect RDH12 expression, membranes were blocked in 5% non-fat milk, then incubated with an HRP-conjugated mouse monoclonal anti-polyHistidine antibody (Sigma, A7058, RRID: AB_258326; 1:2500) for 2 hours at room temperature with mild rocking. Signals were visualized via enhanced chemiluminescence (ECL) using a GE ChemiDoc imaging system. GAPDH was used as an internal loading control (10494-1-AP from Proteintech, RRID: AB_2263076; 1:10,000).

#### Multi-omics Sample Preparation

Simultaneous extraction of lipids and proteins from cell pellets was carried out using a monophasic solvent system composed of *n*-butanol:acetonitrile:water (3:1:1, v/v/v), as previously described (44, 45). For lipid extraction, 200 μL of the monophasic solvent was added to each cell pellet, followed by vortexing for 10 s and incubation at 4 °C for 15 min. Samples were then centrifuged at 15,000 × *g* for 15 min at 4 °C. The resulting supernatant, containing the lipid fraction, was carefully transferred to a fresh vial and dried using a SpeedVac vacuum concentrator (MPF Facility).

#### Proteomic Extraction

The remaining protein pellet was resuspended in 300 μL of lysis buffer containing 50 mM ammonium bicarbonate (AmBic), 12 mM *N*-lauroylsarcosinate, 0.5% sodium deoxycholate, and 1× Halt™ Protease Inhibitor Cocktail (Thermo Fisher Scientific, Waltham, MA). To shear DNA, we performed probe sonication with samples on ice (5 s on /5 s off cycles, total 45 s, 40% amplitude; Fisher Scientific, Hampton, NH). Lysates were subsequently clarified by centrifugation at 20,000 × *g* for 30 min at 4 °C. Protein reduction was achieved by adding 6.6 μL of 100 mM dithiothreitol (DTT) and incubating at 50 °C. Alkylation was performed by adding 10 μL of 100 mM iodoacetamide (IAA) and incubating at room temperature in the dark for 30 min. The reaction was quenched with an additional 6.6 μL of 100 mM DTT.

#### SP3 Clean-up and Digestion

For proteomic clean-up, 100 μg of protein per sample was processed using the Single-Pot Solid-Phase-enhanced Sample Preparation (SP3) method. A bead-to-protein ratio of 7:1 was employed, using a 1:1 mixture of hydrophobic and hydrophilic Sera-Mag SpeedBeads Carboxyl Magnetic Beads (Cytiva Life Sciences, Marlborough, MA; Cat# 45153105050250 and 6515210505250), prepared at 20 μg/μL. Proteolytic digestion was carried out in 100 mM AmBic (pH 8.4) using Trypsin/Lys-C Mix, mass spectrometry grade (Promega, Madison, WI), at a 25:1 protein-to-enzyme ratio, and incubated overnight at 37 °C. After digestion, the peptide-containing supernatant was transferred to a new Lo-Bind Eppendorf tube. Peptides were acidified with formic acid to a final concentration of 0.1% (v/v) prior to desalting.

#### Desalting

Desalting of digested peptides was performed using C18 solid-phase extraction (SPE) tips (Affinisep, Normandy, France). Tips were conditioned first by sequentially passing through 100 μL of ACN, followed by 100 μL of 20% ACN in 0.1% FA, with each step centrifuged at 200 × *g* for 2 minutes. The tips were then equilibrated twice with 150 μL of 0.1% FA in water, each centrifuged at 200 × *g* for 3 minutes. Acidified peptide samples (final concentration 0.1% FA) were loaded onto the tips and centrifuged at 200 × *g* for 3 minutes. The flow-through was reloaded to ensure maximal binding, then centrifuged again under the same conditions. Bound peptides were washed three times with 150 μL of 0.1% FA in water, each wash followed by centrifugation at 200 × *g* for 3 minutes. Peptides were eluted sequentially with 60 μL of 50% ACN with 0.1% FA and 60 μL of 80% ACN with 0.1% FA, each step centrifuged at 200 × *g* for 2 minutes. The desalted peptide solutions were transferred to Lo-Bind Eppendorf tubes, dried using a refrigerated vacuum concentrator (Labconco Cold SpeedVac, Kansas City, MO), and stored at −80 °C until reconstitution for LC–MS analysis.

#### Proteomics Data Acquisition on Exploris 480

Peptides were reconstituted in 98:2 H_2_O:ACN containing 0.1% FA. Peptides were analyzed via a Vanquish Neo UHPLC coupled to a Thermo Orbitrap Exploris 480 (Thermo Fisher Scientific, Waltham, MA). Chromatographic separation was performed on a PepMap Neo C18 column (75 µm x 50 cm, 2 µm particle size, double nanoViper, Thermo Fisher) at 50°C. Mobile phase A consisted of water with 0.1% FA, and mobile phase B consisted of acetonitrile with 0.1% FA. Peptides were separated over a 90-min gradient from 4-36% B at a flow rate of 0.3 μL/min with a total acquisition time of 106 mins (0.4-4% B 0-1 min,4-24% B 1-81 min, 24-36%B 81-91 min, 36-80% B 91-96 mins, 80% B 96-106 min, followed by column equilibration to 0.4% B). A nanospray ionization source with a nano-bore stainless steel emitter (150 µm x 30 µm) at a spray voltage of 2.0 kV and an ion transfer tube at 280 °C.

We used variable staggered-window data-independent acquisition (DIA) in positive ionization mode, with an MS1 scan at 120,000 mass resolution at *m/z* 200 from 374 to 1,200 *m/z*, a normalized AGC target of 300% (absolute AGC target of 3×10^6^), and an RF lens of 50%. DIA scans were acquired at 22,500 mass resolution at *m/z* 200 with a scan range of 145-1450 *m/z*, a max injection time of 38 ms, a normalized AGC target of 1000% (absolute AGC target of 1×10^6^), RF of 50%, and a normalized higher-energy collisional dissociation energy of 30%. We used 52 custom-defined, variable-width DIA windows with 0.5 Th overlap, with widths of either 12 m/z or 24 m/z depending on precursor density in HEK293T tryptic digests. We optimized the precise window boundaries and overlap parameters based on empirical peptide distribution, as detailed in **SI Table-1**.

#### Protein Extraction and S-Trap Sample Preparation for LC–MS/MS on timsTOF HT

HEK293T (WT &RDH12) cells were lysed in RIPA buffer supplemented with a protease inhibitor cocktail. Cell lysates were homogenized using a Precellys bead homogenizer and centrifuged at 14,000 × *g* for 20 min at 4 °C. We collected the clarified supernatants and determined protein concentrations using the Pierce™ BCA Protein Assay Kit (Thermo Fisher Scientific, Cat. No. 23227) according to the manufacturer’s instructions. For proteomic sample preparation, we denatured 100 µg total protein in 8 M urea for 10 min at room temperature, reduced it with 5 mM dithiothreitol (DTT) at 60 °C for 1 h, and alkylated it with 15 mM iodoacetamide (IAA) for 30 min at room temperature in the dark. Protein samples were acidified with phosphoric acid to a final concentration of approximately 2.5% and mixed with S-Trap protein binding buffer. The acidified protein mixture was loaded onto S-Trap Micro spin columns (Protifi, Cat. No. C02-micro-80). After binding, the columns were washed three times with chloroform/methanol, then with S-Trap binding/wash buffer to remove detergents and other contaminants. Proteins were digested on-column with 2 µg of Trypsin/Lys-C Mix (Thermo Fisher Scientific, Cat. No. A41007) overnight at 37 °C. Peptides were sequentially eluted with 50 mM triethylammonium bicarbonate (TEAB), 0.2% formic acid in water, and 50% acetonitrile. The eluates were pooled and dried to completion in a SpeedVac concentrator.

Peptides were desalted using Pierce™ C18 Spin Columns (Thermo Fisher Scientific, Cat. No. 89870) according to the manufacturer’s protocol. Desalted peptides were eluted with acetonitrile containing formic acid, dried by vacuum centrifugation, and quantified using the Pierce™ Quantitative Colorimetric Peptide Assay (Thermo Fisher Scientific, Cat. No. 23275) before loading.

#### Proteomics Data Acquisition on timsTOF HT

Peptides were separated on a Bruker nanoElute nano-flow UHPLC system operated in two-column (trap-and-elute) mode and analyzed on a Bruker timsTOF HT trapped ion-mobility quadrupole time-of-flight mass spectrometer equipped with a CaptiveSpray nano-electrospray ion source (timsControl acquisition software v6.0.8; HyStar v6.3.1.8). Peptide (∼200 ng) from each sample was injected (2 µL, µL-pickup partial-loop injection) and loaded onto a trap cartridge (Thermo PepMap Neo Trap Cartridge, 5 mm × 300 µm i.d., 5 µm C_18_ particles) before separation on a Bruker PepSep Classic analytical column (25 cm × 150 µm i.d., 1.5 µm C_18_ particles) held at 50 °C. Peptides were eluted at a constant flow rate of 0.35 µL/min over a 70 min method using mobile phases [A: 0.1% (v/v) formic acid in water] and [B: 0.1% (v/v) formic acid in acetonitrile]. The gradient ramped from 3% to 26% B over 50 min, to 32% B at 60 min, then to 95% B at 60.5 min, and was held at 95% B until 70 min.

The mass spectrometer was operated in positive-ion dia-PASEF mode. The CaptiveSpray source was operated with a capillary voltage of 1600 V, end-plate offset of -500 V, dry gas of 3.0 L/min and a dry-gas temperature of 180 °C. Mass spectra were recorded over m/z 100-1700 and an ion-mobility (1/K0) range of 0.70-1.30 V·s/cm². The TIMS analyzer was run with equal accumulation and ramp times of 100 ms each (100% duty cycle), corresponding to 944 TIMS scans per frame. The dia-PASEF isolation scheme comprised 8 window groups and 38 total isolation windows of 25 Th width, tiled across the *m*/*z* range 228-1178 and staggered in ion-mobility space so that each dia-PASEF frame sampled multiple precursor windows. The collision energy was ramped linearly as a function of ion mobility from 20 eV at 1/K0 = 0.6 V·s/cm² to 59 eV at 1/K0 = 1.6 V·s/cm² (≈24-46 eV across the acquired mobility range), with a collision-cell RF of 1500 Vpp.

#### Lipidomics Data Acquisition

Lipid extracts were reconstituted in a solvent mixture consisting of acetonitrile:methanol:300 mM ammonium acetate (3:6.65:0.35, v/v). We performed discovery-based untargeted lipidomic analysis using multiple reaction monitoring (MRM) profiling (46). In brief, the MRM profiling was across 11 lipid classes and MRM transitions were selected to detect class-diagnostic product ions for glycerophospholipids, including headgroup-associated fragments for phosphatidylcholine, phosphatidylethanolamine, and other phospholipid classes. For diacylglycerols and triacylglycerols, transitions were based on neutral losses corresponding to fatty acyl chains. A 10 μL aliquot of each sample was directly injected into the electrospray ionization (ESI) source (G1377A) of an Agilent 6410 triple quadrupole mass spectrometer (Agilent Technologies, Santa Clara, CA, USA) via flow injection. A capillary pump was connected to the autosampler and operated at a flow rate of 8 μL/min and pressure of 150 bar. The capillary voltage was 5 kV, and the gas flow was 5.1 L/min at 300 °C. Raw MS data were collected as Agilent .d folders.

#### Data processing and analysis

Proteomic: Peptide identification, protein inference, and label-free quantification were performed in DIA-NN (47) version 2.1.0 for the Exploris 480 data and version 2.2.0 for the timsTOF HT data, using a library-free (in silico predicted library) workflow. The predicted spectral library was generated with DIA-NN (version 1.9.1) by in silico tryptic digestion of the canonical human reference proteome (UniProt UP000005640_9606, downloaded 26 May 2024; 20,654 entries) concatenated with the cRAP contaminant database (camprotR build of the 2019-04-01 cRAP release), 20,736 protein entries in total, with Trypsin/P specificity, up to one missed cleavage, peptide length 7-30, precursor *m*/*z* 300-1,800, precursor charge 1-4, *N*-terminal methionine excision, carbamidomethylation of cysteine as a fixed modification, and oxidation of methionine and protein *N*-terminal acetylation as variable modifications (maximum one per peptide). Fragment ion *m*/*z*, relative intensity, retention time, and ion mobility were predicted for every precursor, retaining up to 12 fragment ions per precursor, yielding 4,075,657 target precursors. Decoy precursors were generated by DIA-NN at a 1:1 ratio by residue mutation of each target sequence and searched together with the targets; precursor and protein-group q-values were estimated from the target-decoy score distributions of DIA-NN’s neural-network classifier.

Raw files were searched directly. Mass accuracies were optimized on the first run and applied to all runs (MS1 9 ppm and MS2 20 ppm for the Exploris 480 data; 20 ppm for both for the timsTOF HT data). Precursor (MS1) signal was extracted for every library precursor and contributed both to the identification score (MS1 profile correlation with the fragment traces) and to quantification. Predicted retention times were aligned to each run by DIA-NN’s non-linear calibration on confidently identified precursors; the retention-time extraction window was set automatically at approximately 6 min in the first pass and 2 min in the second pass, with a scan-window radius of 7 cycles (Exploris) or 15 cycles (timsTOF). For the timsTOF data, predicted ion mobilities were calibrated per run in the same way, with 1/K₀ extraction windows of approximately 0.045 and 0.01 V·s/cm² in the first and second pass, respectively. Match-between-runs was enabled: a first pass over all runs of an experiment produced an empirical library at 1% precursor FDR (135,883 precursors, 106,809 peptides, 9,214 protein groups for the 27 Exploris runs; 109,814 precursors and 9,784 protein groups for the six timsTOF runs), which was used to re-analyze all runs in a second pass. Peptidoform scoring was enabled. Protein inference used proteotypic (unique) peptides only, and protein groups were quantified by the MaxLFQ algorithm as implemented in DIA-NN with retention-time-dependent cross-run normalization with cRAP entries excluded from normalization and quantification. Results were filtered to run-specific precursor q-value ≤ 0.01, run-specific protein-group q-value ≤ 0.05, global precursor q-value ≤ 0.01, global protein-group q-value ≤ 0.01, channel q-value ≤ 0.05, and proteotypic precursors only, and contaminant protein groups were removed, yielding 8,116 protein groups across the 27 Exploris runs and 8,389 across the six timsTOF runs. The number of distinct peptide sequences and precursors supporting each protein group, and the number of proteotypic peptides for that group in the refined library, are given in SI Tables 2-4. Log_2_-transformed MaxLFQ protein quantities were exported (Parquet) for statistical analysis in R (SI Table-5). For each contrast, protein groups were classified by the missingness of the two replicates compared. Where neither condition contained an observed value in any replicate, the comparison rests on imputed values and was excluded from testing. Where only one conditioned lacked observed values, the protein was tested but is flagged as a presence/absence call, and its fold change should be read as directional rather than quantitative (**SI Table-6**). We then adjusted the p-values using the Benjamini-Hochberg method to control the false discovery rate.

Single-peptide protein groups. Protein groups were retained irrespective of the number of supporting peptides. Each was annotated with the number of distinct proteotypic peptides passing the q-value filters and, for single-peptide groups (479 of 8,049), assigned a supporting tier from the median DIA-NN Evidence score across identified runs (tier B, ≥4; tier C, <4) (**SI Table-2**). To provide chromatographic evidence, the 27 raw files were re-search in DIA-NN 2.1.0 against the refined spectral library from the primary search with identical parameters and extracted chromatograms exported for all precursors (--xic 90). Precursor and fragment traces for every single-peptide protein group across all 27 runs are provided in SI Data 1, and those for proteins discussed individually in the text as SI Data 2. Proteins whose chromatograms lacked coeluting fragment ions were not used as individual evidence and are excluded from heatmaps. Single-peptide groups that passed inspection are marked with an asterisk.

Lipidomics: The raw MS data were processed using an in-house script that exports each MRM transition and its absolute ion intensity to Microsoft Excel. Lipid identification was performed by extracting absolute ion intensities for all monitored MRM transitions and filtering signals observed at least 1.3 times higher than in a blank sample SI Table-7-8. These results represent tentative lipid identifications from discovery-mode MRM profiling and have not been confirmed by targeted LC-MS/MS analysis using internal standards. Lipid annotations were parsed from candidate lipid names derived from the LIPID MAPS database and reported using lipid shorthand nomenclature consistent with current lipidomics annotation guidelines. We handled acute and recovery lipid datasets separately because samples were extracted and collected on different dates, and PCA of the combined datasets showed high variance between them. We normalized the data using variance-stabilizing normalization (“vsn”) in the Bioconductor package DEP SI Table-9-10.

Gene ontology (GO) enrichment analysis was performed in R using clusterProfiler::enrichGO() on proteins significantly increased or decreased in each contrast, defined by an absolute fold change of at least 2 and an adjusted p value < 0.01. We simplified GO biological process terms using clusterProfiler::simplify(), with semantic similarity information generated using GOSemSim::godata() for human biological process terms. Pathway enrichment was performed using Reactome pathway annotations with ReactomePA::enrichPathway().

All statistical analyses and visualizations were performed in R. Code used for data analysis is available at github.com/drown-lab/atRAL-RDH12-HEK and Zenodo (DOI 10.5281/zenodo.22943041).

## Results

### Stable Expression of RDH12 Alleviates atRAL-Mediated Cytotoxicity via Reductive Clearance at the Endoplasmic Reticulum

To investigate intracellular defense mechanisms against reactive retinaldehyde stress, we first established a stable HEK293T expression system for the human short-chain dehydrogenase/reductase RDH12 using a GFP-tagged construct (RDH12), alongside an empty-vector control line expressing unconjugated GFP (WT). Confocal microscopy verified the subcellular localization of RDH12 in HEK293T cells. WT cells expressing free GFP showed a diffuse, whole-cell distribution. In contrast, the RDH12-GFP fusion protein exhibited a distinct reticular cytoplasmic signal that co-localized with ER-Tracker Red, confirming RDH12 localization to the endoplasmic reticulum membrane (**Figure 2A**). Western blot analysis validated the robust expression of the RDH12-GFP enzyme at its expected molecular weight (62 kDa) (**Figure 2C**). To assess whether overexpression of this RDH12 construct caused broader proteome changes, we performed bottom-up proteomic analysis under basal conditions. RDH12-expressing and WT cells showed highly similar proteomic profiles, sharing over 99% of detected protein groups and a Pearson correlation coefficient of 0.99 in protein abundances, with RDH12 detected exclusively in RDH12-expressing cells . We validated ER stress in RDH12-expressing cells and found no detectable stress (**Figure S1 A-D**).Next, we verified that the localized enzyme maintained high-velocity catalytic fidelity in intact cells by tracking the metabolic flux of an acute 5 µM all-*trans*-retinal (atRAL) challenge. After 4 h of exposure, control cells converted a modest portion of the aldehyde to retinol (3.00±0.10 nmol/mg), while a notable fraction was oxidized to retinoic acid (0.97±0.06 nmol/mg) (**Figure 2B**). Conversely, cells stably expressing RDH12 exhibited a nearly 3-fold spike in retinol production (8.90±0.61 nmol/mg), while simultaneously suppressing retinoic acid formation by over 87% (0.12±0.01 nmol/mg). This confirms that ER-resident RDH12 efficiently competes with endogenous aldehyde dehydrogenases, shunting reactive retinal toward a reductive pathway.

**Figure 2.**
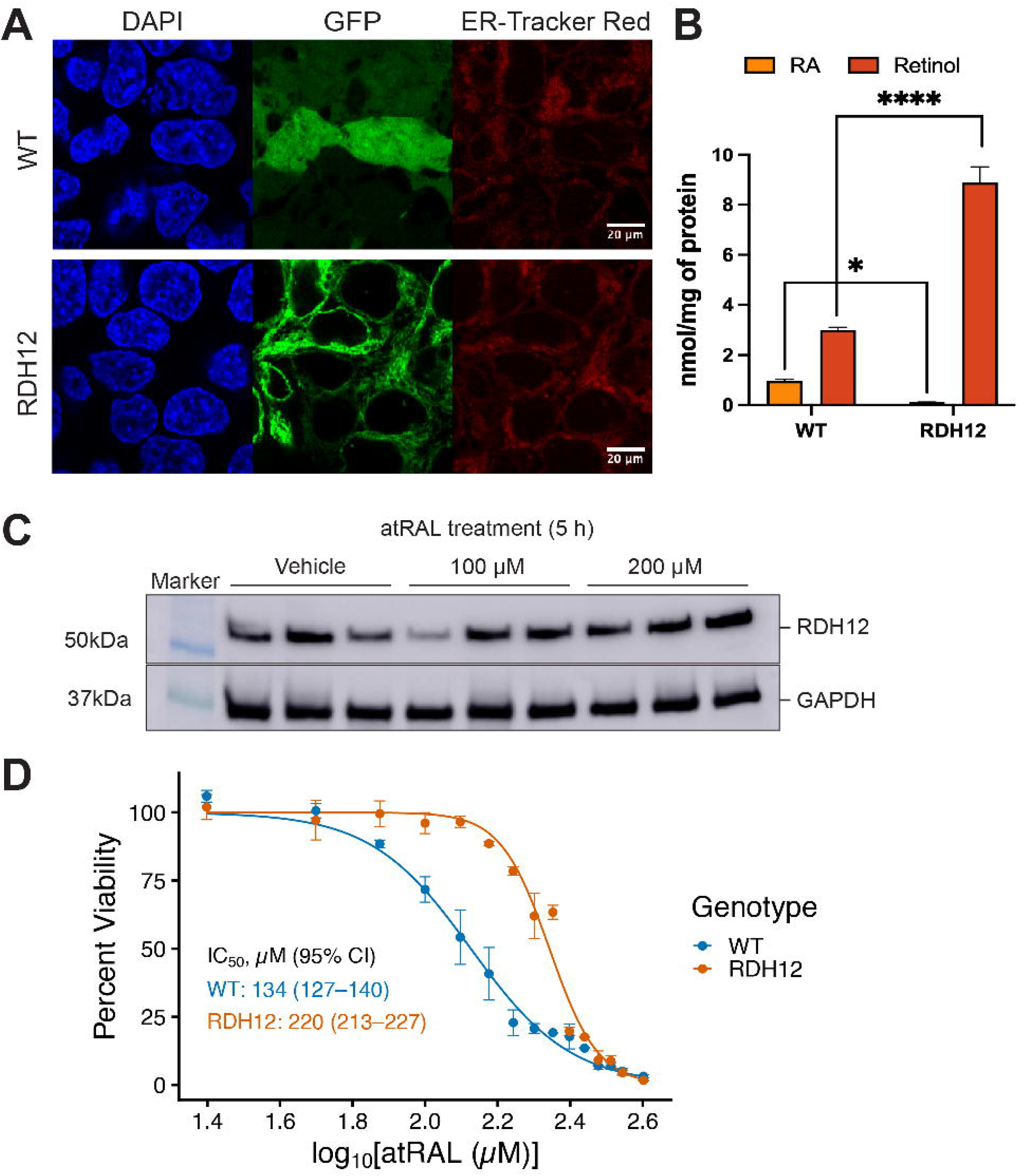
Characterization, expression, and cytoprotective activity of RDH12 under atRAL stress. *A*, Representative confocal microscopy images of WT and RDH12-GFP-expressing HEK293T cells. Cells were counterstained with DAPI (blue, nuclei) and ER-Tracker Red (red, endoplasmic reticulum). The RDH12-GFP construct (green) demonstrates a characteristic reticular distribution that colocalizes with the ER marker, confirming successful endoplasmic reticulum targeting, whereas the WT control exhibits a diffuse cytosolic GFP distribution. Scale bars: 20 µm. *B,* Functional retinal reductase activity of cell lysates following an acute challenge with 5 µM atRAL for 4 h. The accumulation of retinol (ROL) and retinoic Acid (RA) in the medium was quantified via HPLC and normalized to total cellular protein (n=3; mean ±SD; Statistical significance was determined using a Two-way ANOVA followed by Šídák’s multiple comparisons test \**p*<0.05, \*\*\*\**p*<0.0001). *C*, Immunoblot verification of recombinant RDH12-GFP expression using anti-His antibody in cells treated with different concentrations of atRAL (Vehicle, 100 µM, and 200 µM) for 5 hours. GAPDH is shown as a loading control to confirm equal protein distribution across all experimental conditions. *D*, Dose-response cell viability curves of control and RDH12-expressing cells exposed to varying concentrations of atRAL (0-400 µM) for 5 h, measured via CellTiter-Glo luminescence and normalized to untreated baselines (n=3).

Finally, we tested whether localized metabolic clearance directly shields cells from aldehyde-driven cell death. Over a 5 h treatment window spanning a broad concentration range (0-400 µM atRAL), control cells underwent a rapid, dose-dependent collapse in viability, yielding a calculated IC_50_ of 134 µM (95% CI: 127 to 140 µM) (**Figure 2D**). In contrast, cells expressing RDH12 demonstrated a prominent rightward survival shift, resisting atRAL-mediated toxicity with an IC_50_ of 220 µM (95% CI: 213 to 227 µM). Together, these data demonstrate that the retinal reductase activity of ER-localized RDH12 rescues cells from atRAL cytotoxicity.

### Effect of Acute atRAL-induced toxicity on the proteome in RDH12-expressing HEK293T Cells

To assess the effect of atRAL-induced toxicity on the proteome, we induced acute aldehyde toxicity in a HEK293T cell line overexpressing RDH12 by treating cells with atRAL for 5 h at 100 µM and 200 µM, with a vehicle (Veh) treatment of ethanol (**Figure 1A**). We applied a monophasic extraction protocol to isolate lipids and proteins from these samples for pair analysis (*45*). We performed untargeted bottom-up proteomics using a data-independent acquisition strategy and quantified 8,049 protein groups across the acute and recovery datasets (SI Table 5 and **Figure S2A**). Stable expression of RDH12 remained high across treatments (**Figure S3)**. To assess data quality before imputation and differential abundance testing, we examined protein detection overlap and replicate-level variability (**Figure S2B-C**). Principal component analysis of pre-imputation protein intensities showed clear separation of the acute RDH12 vehicle, 100 µM, and 200 µM atRAL samples, indicating that acute atRAL exposure produced dose-dependent proteomic changes (**Figure S2D**). In the recovery dataset, PCA separated RDH12-expressing cells from WT cells, supporting genotype-dependent differences in the post-atRAL recovery proteome (**Figure S2E**). Because missing values are common in discovery proteomics (48), we evaluated missingness patterns and imputation behavior before differential abundance testing. Missing proteins were enriched among lower-abundance proteins, and post-imputation intensity distributions remained comparable to the filtered pre-imputation data (**Figure S4**). We identified differentially abundant protein groups across key contrasts using limma analysis). Acute atRAL exposure induced extensive proteome remodeling. Relative to the vehicle, 100 µM atRAL increased 534 proteins and decreased 1,831 proteins (**Figure 3A**), whereas 200 µM atRAL increased 666 proteins and decreased 813 proteins (**Figure 3B**), based on thresholds of adjusted p-values < 0.01 and fold change > 2. Detection depth differed across sample groups, and the acute vehicle group was the least deeply sampled: 5,984 protein groups were quantified in all three replicates, compared with 6,358 to 7,514 in the other eight groups, and 893 proteins were undetected in all three acute vehicle replicates, compared with 194 to 418 elsewhere (**Figure S2A**). As a consequence, 359 of the 534 increases at 100 µM and 452 of the 666 increases at 200 µM involve proteins not detected in any vehicle replicates, so their reported fold changes are determined by the imputation model rather than by direct measurement. Decreases are far less affected, with only 3 to 5 percent arising from non-detection in the treated group. Fold-change magnitudes for proteins with a missing reference condition should therefore be read as directional. The missingness class of every protein in every group is reported in SI Table 6.

**Figure 3.**
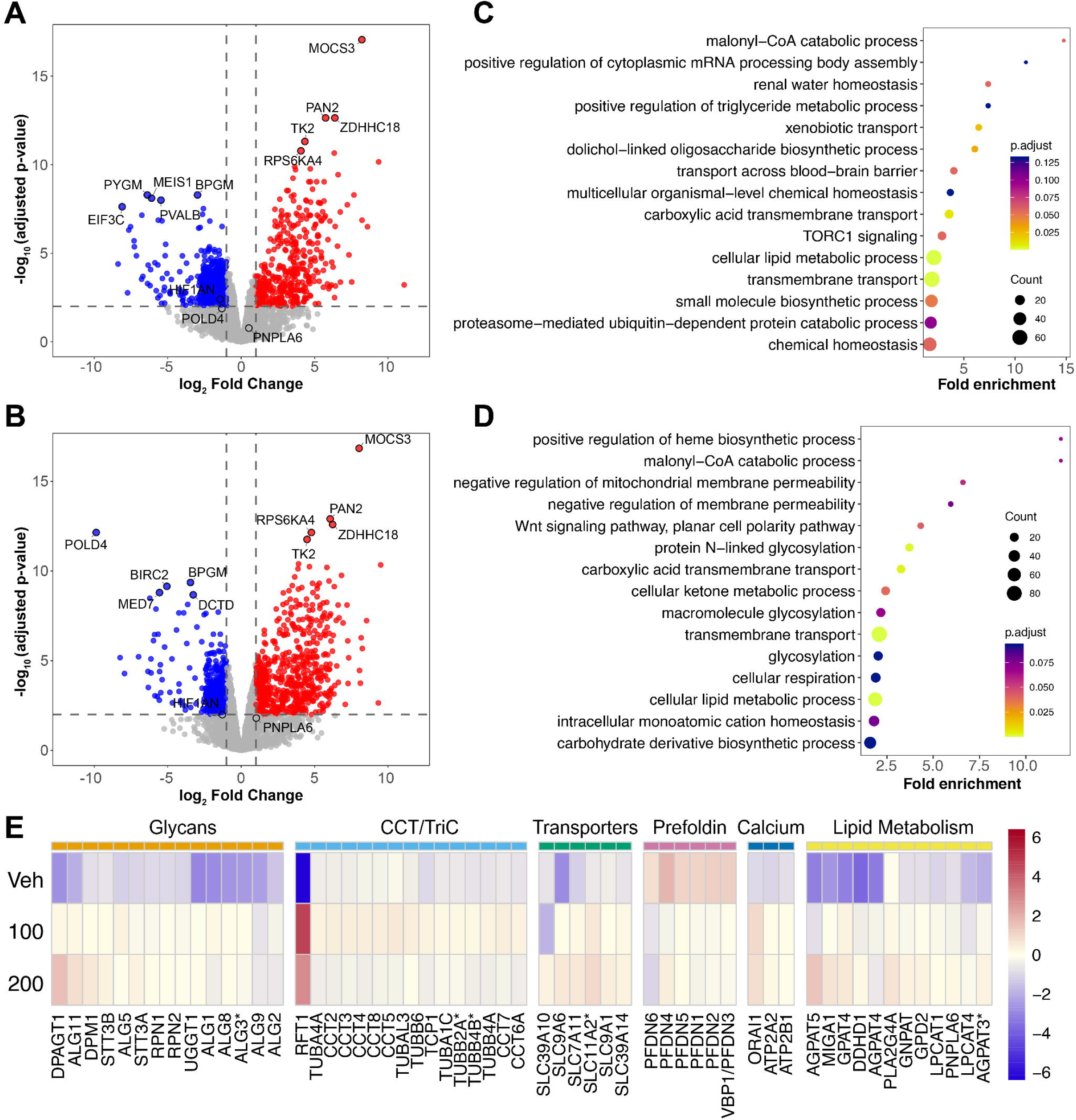
Acute atRAL-toxicity proteome response. Volcano plots showing global proteome changes in response to acute atRAL exposure at (A) 100 µM and (B) 200 µM. Vertical dashed lines indicate ±2-fold change, and the horizontal dashed line indicates the BH-adjusted p-value cutoff of 0.01. Proteins highlighted in red are enriched upon atRAL exposure, whereas proteins highlighted in blue are enriched in the vehicle. (C) Biological process GO enriched terms enriched upon atRAL (C) 100 µM and (D) 200 µM. (E) The heatmap shows selected proteins of interest from GO terms of interest that met a significance threshold of fold change ±1.4 and BH-adjusted p < 0.1. Colors represent protein-wise mean-centered log_2_ intensities within each condition. Asterisks mark protein groups identified by a single proteotypic peptide. Their extracted ion chromatograms are provided in SI Data 2.

Among the proteins detected only after treatment were MOCS3, PAN2, and ZDHHC18 (**Figure 3A, B**). MOCS3 is involved in molybdenum cofactor biosynthesis and tRNA thiolation (49), and ZDHHC18 is a palmitoyltransferase linked to membrane-associated protein regulation (50). PAN2, also called RDH14, is an NADPH-dependent microsomal enzyme in the short-chain dehydrogenase/reductase superfamily that efficiently reduces all-trans-retinaldehyde to all-trans-retinol and is ubiquitously expressed in human tissues (51). Its selective detection after atRAL treatment indicates that PAN2 may be recruited as a compensatory retinal reductase to support RDH12-mediated aldehyde clearance. In contrast, HIF1AN decreased at both atRAL concentrations (log_2_FC = -1.4 at 100 µM and -1.3 at 200 µM), consistent with altered protein abundance linked to hypoxia-associated signaling (52). Observed in vehicle-treated cells, POLD4 was not detected after 200 µM atRAL treatment, which may reflect disruption of DNA replication processes, as its degradation is required to inhibit fork progression and cell survival (53). Together, these individual protein changes point to altered RNA processing, sulfur metabolism, regulation of membrane-associated proteins, and genome maintenance during acute atRAL exposure.

To identify biological processes associated with the acute response to atRAL, we performed Gene Ontology (GO) enrichment analysis using proteins significantly increased or decreased in each contrast and all quantified proteins as the background set. Proteins with increased abundance following atRAL exposure were enriched for terms related to N-glycan biosynthesis, solute carrier-mediated transport, lipid metabolic processes, and membrane-associated stress responses (**Figure 3C, D**). In contrast, proteins more abundant in vehicle-treated cells were enriched for translation, lysosome, vesicle, membrane regulation, gene expression, and RNA- and DNA-associated processes (**Figure S5**). Together, these results indicate that acute atRAL exposure is associated with the regulation of biological processes involved in ER protein processing, ion and metabolite transport, cytoskeletal proteostasis, and lipid metabolism.

With attention drawn to these processes and pathways, heatmaps of individual proteins contributing to enriched biological processes and pathways were constructed (**Figure 3E**). Multiple proteins involved in N-linked glycosylation and ER protein processing, including ALG1, ALG11, DPAGT1, STT3A, STT3B, RPN1, RPN2, and UGGT1, were increased following atRAL exposure. This observation is consistent with the increased demand for the ER secretory pathway and protein quality control machinery during acute stress (*54*). Ion and metabolite transport proteins also contributed to the atRAL-induced proteomic response (**Figure 3E**). Increased abundance of SLC39A14 and SLC11A2 is consistent with altered iron-associated transport pathways (55, 56). The detection of SLC7A11 only after atRAL exposure may reflect an increased demand for cystine import to support glutathione synthesis and antioxidant defense (*57, 58*). In parallel, increased ORAI1, ATP2A2, and ATP2B1 are consistent with remodeling of calcium-handling proteins at the plasma membrane and ER-associated compartments (*59, 60*). These changes suggest that acute atRAL toxicity is associated with coordinated changes in ionic, redox, and proteome homeostasis.

Another feature of the heatmap was the increased abundance of multiple subunits of the chaperonin-containing TCP-1 complex (CCT/TRiC), along with several tubulin proteins, at 100 µM atRAL (**Figure 3E**). This increase was not sustained at 200 µM, where CCT subunit abundance returned to vehicle levels. CCT/TRiC is an ATP-dependent group II chaperonin that folds a substantial fraction of the cytosolic proteome, and chaperone capacity has been proposed as a determinant of photoreceptor viability in retinal dystrophies (*61, 62*). In contrast, several subunits of the prefoldin complex were decreased following atRAL at both doses. Prefoldin is a holdase and ATP-independent co-chaperone that binds to unfolded actin and tubulin and delivers them to CCT/TRiC for folding (*63, 64*). This divergent observation indicates that acute atRAL stress does not uniformly increase all components of the cytoskeletal protein-folding pathway. This is consistent with selective regulation of proteostasis, in which increased CCT/TRiC abundance may reflect an elevated demand for folding or stabilization of protein clients, whereas reduced prefoldin abundance may accompany suppression of new protein synthesis or impaired client delivery to the CCT/TRiC folding chamber (*65–67*). Overall, the acute proteomic data show that atRAL exposure alters proteins associated with ER protein processing, cytoskeletal folding, transporter processes, and translation-related processes. However, we found no clear signatures of upregulation in unfolded protein response pathways.

### Acute atRAL exposure alters lipid species linked to membrane remodeling

For lipid analysis following multiomics sample preparation, we used discovery-based MRM profiling (*46*) to measure several lipid species. We quantified 226 lipid species in the acute dataset and 325 lipid species in the recovery dataset across 11 lipid classes shown in **Figure 4A** and SI Tables 7 and 8, including phosphatidylcholine (PC), phosphatidylethanolamine (PE), phosphatidylinositol (PI), phosphatidylserine (PS), phosphatidylglycerol (PG), ceramides (Cer), sphingomyelin (SM), acyl-carnitines (Car), triacylglycerols (TG), diacylglycerols (DG), and cholesteryl esters (CE). Because lipid assignments were derived from MRM transitions that could correspond to multiple isobars, we reported individual lipid species as putative annotations from LIPID MAPS. To assess data quality and differential abundance, we examined lipid detection overlap and replicate-level variability (**Figure S6A-B**). Principal component analysis performed on lipid intensities showed clear separation of the acute RDH12 vehicle, 100 µM, and 200 µM atRAL samples (**Figure S6C**), indicating that acute atRAL exposure produced dose-dependent lipidome remodeling. In the recovery dataset, PCA separated RDH12-expressing cells from WT cells (**Figure S6D**), supporting genotype-dependent differences in the post-atRAL recovery lipidome. Differentially abundant lipid species were identified using limma analysis (**SI Tables 9 and 10**).

**Figure 4.**
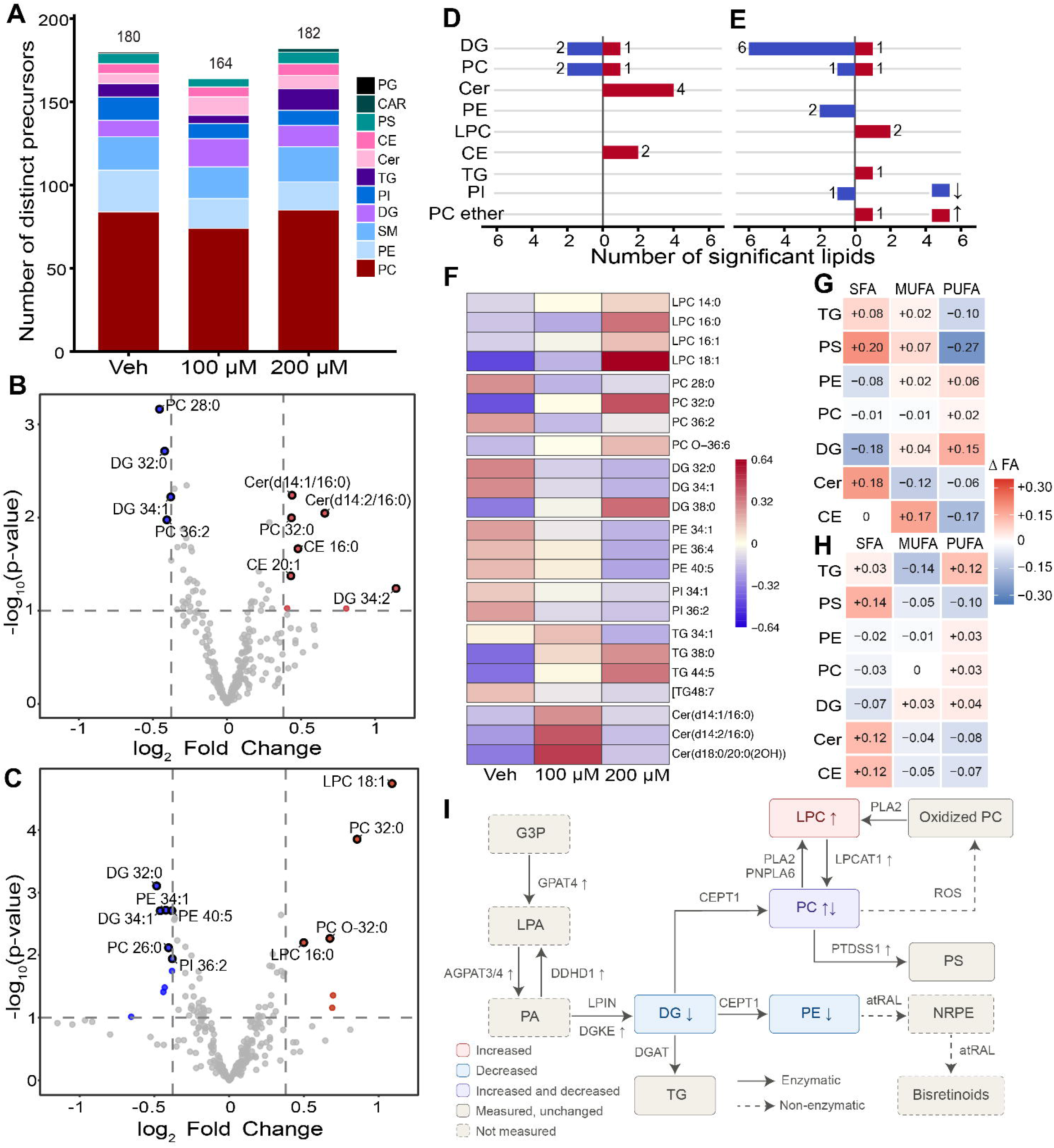
Acute atRAL exposure effects on the lipidome. (A) Number of distinct lipids identified in each lipid class for Vehicle, 100, and 200 µM atRAL-treated groups. Volcano plots showing differentially abundant lipids in (B) 100 µM atRAL vs. Vehicle and (C) 200 µM atRAL vs. Vehicle. Vertical dashed lines indicate ±1.3-fold change, and the horizontal dashed line indicates a nominal p-value cutoff of 0.1. Lipids highlighted in red are enriched upon atRAL exposure, whereas those highlighted in blue are enriched in the vehicle. Lipid class-level directional summary of significant changes in (D) 100 µM atRAL vs. Vehicle and (E) 200 µM atRAL vs. Vehicle comparisons. Red indicates that the number of lipids increased with atRAL treatment, whereas blue indicates that the number of lipids increased in vehicle-treated cells. (F) Heatmap of lipid species of interest across Vehicle, 100 µM, and 200 µM atRAL conditions. Colors represent lipid-wise mean-centered log2 intensities for each condition. The heatmap shows changes in double-bond equivalent (DBE) composition of distinct detected lipid precursors at (G) 100 µM and (H) 200 µM atRAL relative to vehicle. Within each lipid class and treatment group, precursors were grouped as SFA (0 DBE), MUFA (1 DBE), or PUFA (≥2 DBE), and we calculated the proportions of identifications for each class. Tile values indicate the change in precursor proportion compared to the vehicle. Red indicates a higher proportion, and blue indicates a lower proportion relative to the vehicle. (I) Model of acute atRAL-induced lipid remodeling, highlighting measured proteomic and lipidomic changes.

In the acute RDH12-expressing lipidome, lipid changes were subtle after atRAL treatment but indicated a biologically relevant response. Volcano plots comparing 100 µM and 200 µM atRAL to the vehicle showed more significant lipid changes after 200 µM atRAL (**Figure 4B, C**). At 200 µM atRAL, LPC 18:1, LPC 16:0, and PC 32:0 levels were increased relative to those in the vehicle (**Figure 4C, F**). LPC species can be generated through phospholipase-mediated PC cleavage and are commonly associated with membrane turnover, membrane damage, and lipid stress signaling (68, 69). Consistent with this observation is upregulation of patatin-like phospholipase, PNPLA6 in acute atRAL, an enzyme that regulates PC hydrolysis and membrane homeostasis in retinal pigment epithelial cells, and has been genetically linked to RDDs such as LCA and Laurence-Moon syndromes (70, 71).

At the lipid class level, phosphatidylethanolamine (PE) species shifted consistently downward at 200 µM, with 24 of 25 detected showing a negative fold change (**Figure 4C, E**). This consistent direction is notable because atRAL condenses with the primary amine of phosphatidylethanolamine to form N-retinylidene-PE. We detected an increased number of DG species following acute atRAL exposure (**Figure 4A**). We also observed individual DG species changed in both directions across the treatment conditions (**Figure 4D, E**). Because DGs serve as intermediates in glycerophospholipid synthesis and as lipid-signaling molecules, this mixed response is consistent with changes in phospholipid metabolism and DG-associated signaling during atRAL stress (*72, 73*).

At 100 µM atRAL, four ceramide and dihydroceramide species, including C16- and C20-containing species, were increased relative to the vehicle (**Figure 4B, F**). This response differed from that observed at 200 µM atRAL, suggesting that ceramide-associated changes may vary across the atRAL concentration range. Ceramides and related sphingolipids participate in stress signaling, autophagy, apoptosis, and ferroptosis-associated pathways; therefore, their altered abundance may represent one component of the broader lipid stress response to retinaldehyde exposure (74–76). At 200 µM atRAL, selected TG species, including TG 44:5 and TG 38:0, were increased, whereas select PE species were decreased (**Figure 4C,F**).

We evaluated whether acute atRAL exposure was associated with differences in the sum composition of detected fatty acyl (FA) chains, as polyunsaturated FA-containing (PUFA) lipids are susceptible to oxidation. We grouped precursor annotations within each lipid class by the double bond equivalent (DBE): saturated fatty acyl species (SFA; 0 DBE), monounsaturated species (MUFA; 1 DBE), and polyunsaturated species (PUFA; ≥2 DBE), and report the fold-change proportion of DBE following acute atRAL treatment relative to vehicle (**Figure 4G, H**). This enables us to evaluate the proportion of DBE within each lipid class. From this analysis, we observed that DGs had a higher proportion of PUFA-containing precursors at 100 µM atRAL, and TGs had a higher proportion of PUFA-containing precursors at 200 µM atRAL relative to Vehicle. Ceramide precursors had a higher proportion of saturated species after atRAL exposure, and the PC DBE composition remained comparatively stable relative to the vehicle. These observations are notable because PUFA-containing phospholipids and glycerolipids are susceptible to lipid peroxidation (68, 77, 78).

Integrated analysis of lipidomic and proteomic datasets was consistent with altered glycerophospholipid-associated pathways during acute atRAL exposure (**Figure 4I**). The lipidomic data revealed significant changes in phospholipids and glycerolipids, including LPC, PC, DG, ceramide, and TGs. In addition, the proteomic dataset showed an increased abundance of enzymes involved in glycerophospholipid metabolism (**Figure 3E**). GPAT4, AGPAT3, AGPAT4, and AGPAT5 participate in the formation of phosphatidic acid, a central intermediate in phospholipid and neutral lipid synthesis (79). ETNK1 (and, more weakly, CHKA) are associated with Kennedy pathway reactions involved in PC and PE synthesis, whereas PTDSS1 supports PS production from existing phospholipids (80, 81). LPCAT1 and LPCAT4 participate in lysophospholipid reacylation, and DGKE links DG phosphorylation to phosphatidic acid production (*82*). Together, these proteomic and lipidomic changes are consistent with the remodeling of glycerophospholipid-associated pathways during acute atRAL exposure.

### Recovery response to atRAL-induced toxicity reveals genotype-specific stress-resolution

We next examined the molecular changes in the recovery response for RDH12-expressing and WT cells treated with vehicle, 100 µM or 200 µM atRAL for 5h, followed by 24 h recovery in atRAL-free medium before proteomic and lipidomic analysis (**Figure 1A**). In RDH12-expressing cells, the proteomic response after recovery from 100 µM atRAL was limited with only 30 increased and 32 decreased proteins, all based on imputed abundances (**Figure 5A**). Proteins with increased abundance after 100 µM atRAL recovery did not yield significantly enriched GO biological process terms. Proteins with increased abundance in vehicle-treated RDH12-expressing cells were enriched for mitochondrial protein localization and import, indicating that selected proteins involved in mitochondrial trafficking remained lower after 100 µM atRAL recovery (**Figure S7A**). This limited response is consistent with the viability data, in which transient exposure to 100 µM atRAL caused minimal cytotoxicity in RDH12-expressing cells.

**Figure 5.**
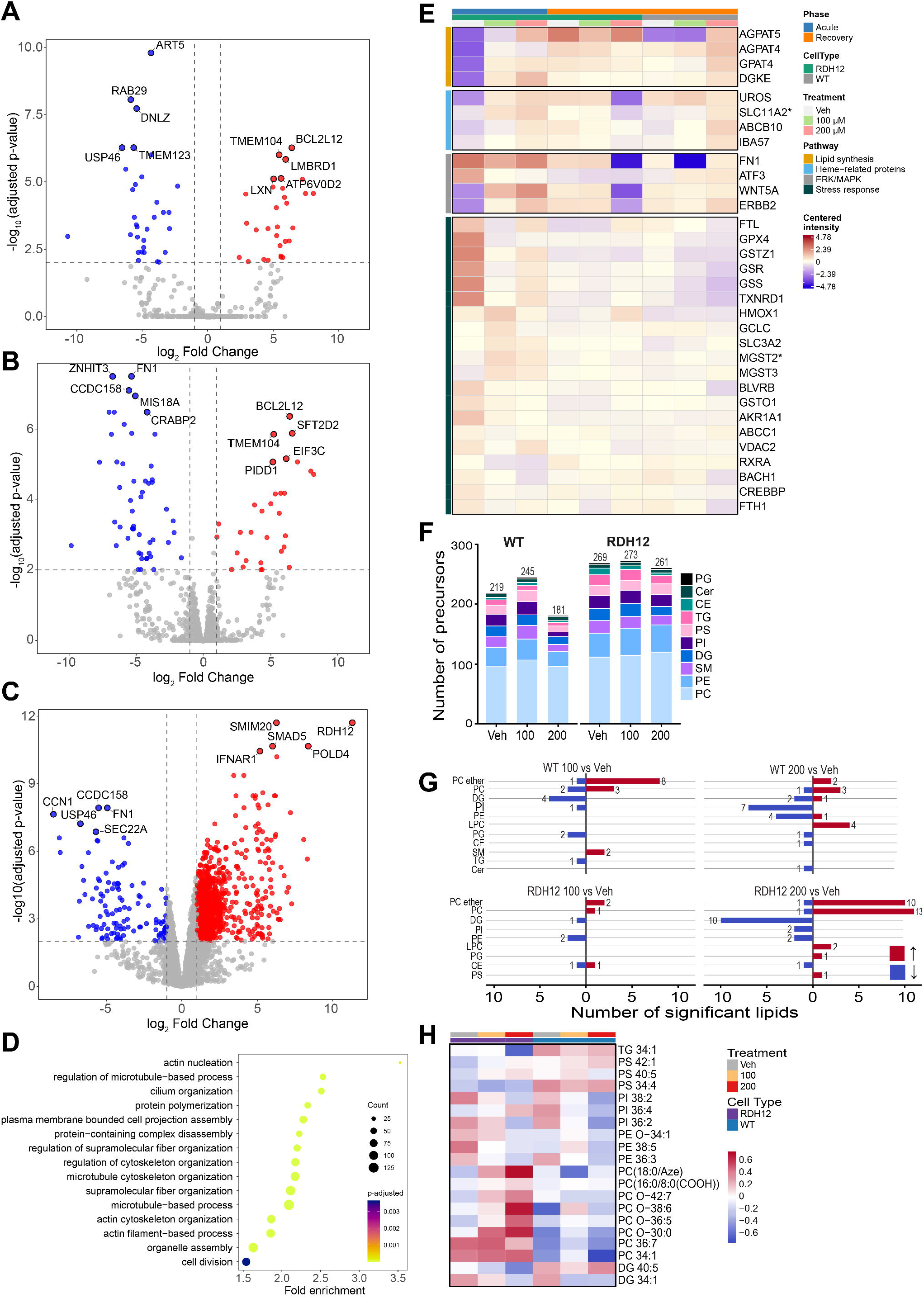
Recovery response to atRAL exposure: Volcano plots showing global proteome changes in response to the recovery response of RDH12-expressing cells at (A) 100 µM and (B) 200 µM. Vertical dashed lines indicate ±2-fold change, and the horizontal dashed line indicates the adjusted p-value cutoff of 0.01. Proteins highlighted in red are enriched upon atRAL exposure, whereas proteins highlighted in blue are enriched in vehicle. (C) Volcano plot of global proteome changes between RDH12-expressing and WT cells at 200 µM atRAL recovery. Vertical dashed lines indicate ±2-fold change, and the horizontal dashed line indicates the adjusted p-value cutoff of 0.01. Proteins highlighted in red are enriched in RDH12-expressing cells, whereas those highlighted in blue are enriched in WT cells. (D) Biological process GO enriched terms from 200 µM atRAL exposure in RDH12-expressing cells relative to WT cells. (E) The heatmap shows selected proteins of interest from GO terms of interest that met a significance threshold of fold change ±1.4 and adjusted p < 0.1. Colors represent protein-wise mean-centered log_2_ intensities within each condition. Asterisk mark protein groups identified by a single proteotypic peptide. Their extracted ion chromatograms are provided in SI Data 2. (F) Number of distinct lipids identified in each lipid class for Vehicle, 100 µM, and 200 µM atRAL treatment in RDH12-expressing and WT cells. (G) Lipid class-level directional summary of significant changes in 100 µM atRAL vs. Vehicle and 200 µM atRAL vs. Vehicle comparisons in each cell type. Red color indicates that the number of lipids increased with atRAL treatment, whereas blue indicates that the number of lipids increased in vehicle-treated cells. (H) Heatmap of lipid species of interest across Vehicle, 100 µM, and 200 µM atRAL conditions in each cell type. Colors represent lipid-wise mean-centered log_2_ intensities within each condition.

Recovery from 200 µM atRAL was associated with a broader proteomic response, with 31 proteins increased and 52 proteins decreased relative to the vehicle (**Figure 5B**). Nearly all of these were presence/absence calls as 28 of the increases were undetected in every vehicle recovery replicate and 49 of the decreases were undetected in every 200 µM-recovery replicate, so their fold changes are set by imputation but are biologically relevant. Proteins with increased abundance after 200 µM atRAL recovery were enriched only for ferroptosis-related GO terms with support from three genes, HMOX1, NFE2L2, and SLC39A7, of which HMOX1 (nine peptides) was quantified in both conditions (**Figures 5E, S7B**). NFE2L2 and SLC39A7 are single-peptide identifications detected only after recovery, and their chromatograms show unique fragment elution (SI Data 2). These proteins are associated with oxidative stress signaling, metal homeostasis, and antioxidant adaptation (83, 84). These findings indicate that, despite atRAL removal, the proteome response of RDH12-expressing cells after recovery from high-dose atRAL exposure is associated with redox regulation, metal handling, and cell-death-associated stress pathways.

After recovery from 200 µM atRAL, proteins with increased abundance in RDH12-expressing cells relative to WT were enriched for microtubule organization and actin cytoskeleton organization (**Figure 5C, D**). In contrast, proteins more abundant in WT cells than in RDH12-expressing cells were associated with ERK/MAPK signaling, fibroblast growth factor receptor signaling, morphogenesis-related processes, and heme biosynthesis pathways (**Figure S7D**). Individual matricellular and extracellular matrix proteins, including CCN1, CCN2 and FN1, were also more abundant in WT cells (**Figure 5C**), although extracellular matrix organization and cell adhesion were not themselves enriched terms (*85, 86*). Among these WT-associated proteins, ATF3, ABCB10, SLC11A2 and IBA57 were quantified in all replicates of both genotypes, whereas CCN1, CCN2, FN1, WNT5A, ERBB2, FGFR2, SPRY2 and UROS were undetected in RDH12-expressing cells, so their fold changes reflect loss of detection (*87–90*). Together, these proteins are consistent with increased extracellular remodeling and growth factor-responsive signaling in WT cells after recovery from high-dose atRAL treatment.

The two genotypes also demonstrated distinct signatures of oxidative stress and metal homeostasis. RDH12-expressing cells showed increased abundance of HMOX1 (**Figure 5E**, log_2_FC 1.28, adjusted p = 2.7 x 10^-5^, quantified in all replicates of both genotypes) and of the NRF2 target SRXN1 (three peptides, detected in all three RDH12 replicates and no WT replicate) (**Figure 6A**), whereas NFE2L2 itself was a single-peptide identification that did not pass chromatographic review (SI Data 2). HMOX1 participates in heme catabolism and is commonly induced during oxidative stress and SRXN1 reduces cysteine sulfinic acids in peroxiredoxins. In contrast, WT cells showed increased abundance of ABCB10, SLC11A2, UROS, and IBA57 (**Figure 5E**), which are linked to heme biosynthesis (**Figure S7D**). Thus, although both genotypes exhibited proteins associated with redox and metal handling after 200 µM atRAL recovery, the RDH12-expressing cells showed a response HMOX1/SRXN1-marked NRF2-target signature, whereas WT cells showed responses centered on heme and cellular metabolism (**Figures 5E, S7**) following atRAL recovery.

**Figure 6.**
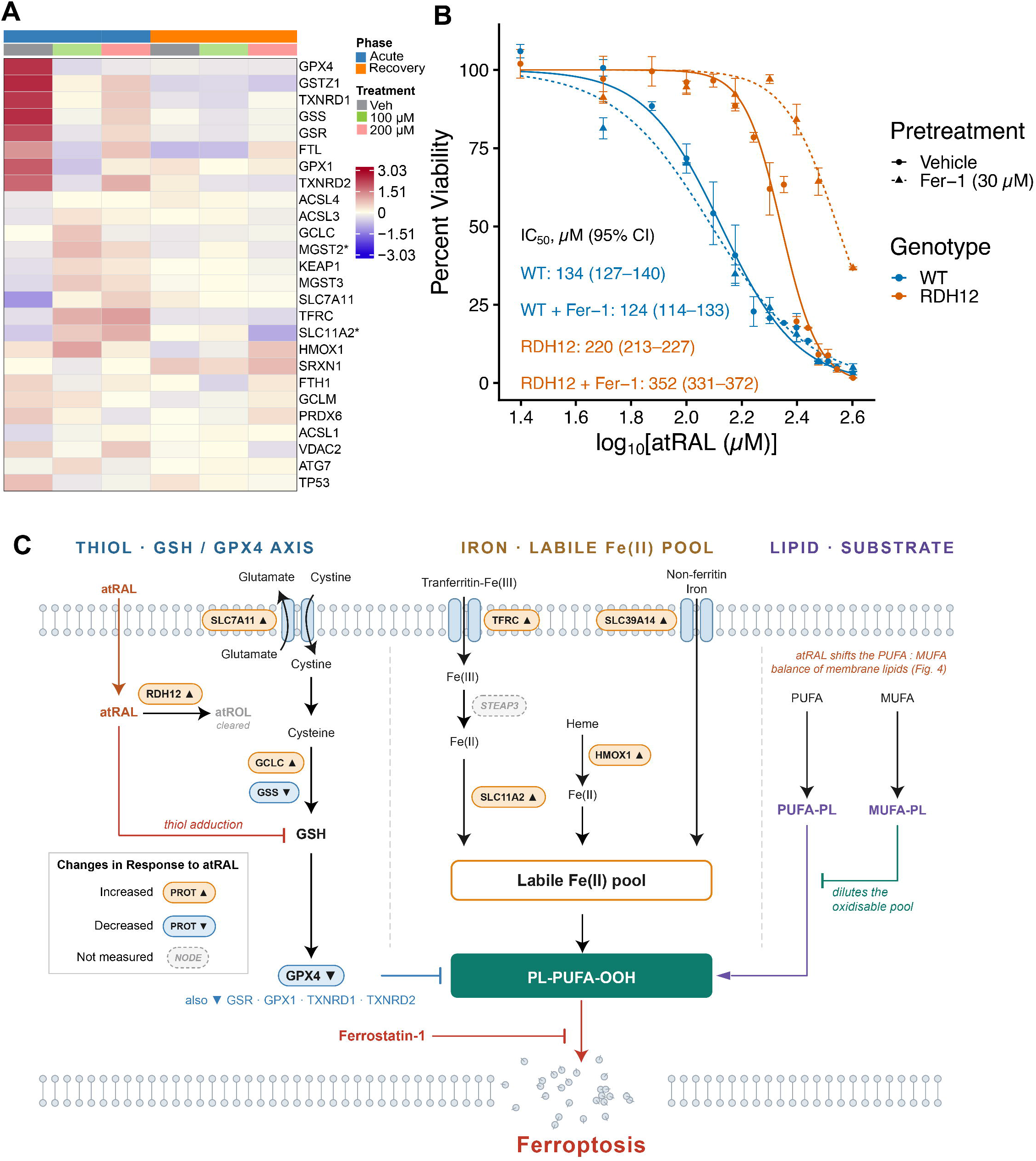
Ferroptosis is an activated form of cell death. (A) The heatmap shows selected KEAP1-NRF2-axis and iron-handling proteins. Per-contrast statistics are in SI Table 2. Colors represent protein-wise mean-centered log_2_ intensities within each condition. Asterisk mark protein groups identified by a single proteotypic peptide. Their extracted ion chromatograms are provided in SI Data 2. (B) Wild-type (WT) and RDH12-expressing cells were pretreated with either vehicle or the ferroptosis inhibitor Ferrostatin-1 (Fer-1; 30 μM) for 24 h and subsequently challenged with increasing concentrations of all-trans-retinal (atRAL; 25-400 μM) for 5 h. We quantified cell viability using the CellTiter-Glo (CTG) luminescent assay and normalized it to untreated controls. Dose-response analysis revealed a modest effect of Fer-1 in WT cells (IC₅₀: 134 μM [95% CI, 127-140 μM] vs. 124 μM [114-133 μM] with Fer-1), whereas Fer-1 markedly increased resistance to atRAL in RDH12-expressing cells, shifting the IC₅₀ from 220 μM (213-227 μM) to 352 μM (331-372 μM). Data are presented as mean ± SEM from three independent biological replicates (N = 3). Each biological replicate was fit separately to give an independent IC_50_, and those values were log_10_-transformed and entered into a two-factor ANOVA with genotype and ferrostatin-1 pretreatment as crossed fixed effects (n = 3 per arm, balanced 2 × 2 × 3 design), followed by Tukey HSD post hoc comparison of the four cell means. (C) The diagram illustrates the competition between atRAL-mediated toxicity and cellular defense mechanisms. Upon entering the cell, atRAL acts as a reactive electrophile that actively suppresses the glutathione (GSH) and thioredoxin-mediated antioxidant response network, thereby unleashing iron-dependent lipid peroxidation. Ferroptosis is promoted by ACSL4-driven incorporation of polyunsaturated fatty acids (PUFAs) into membrane phospholipids (PUFA-PLs). Conversely, cellular survival is mediated by two protective axes: (1) the metabolic clearance of atRAL by RDH12, followed by the DGAT1/2-mediated esterification of retinol into retinyl esters (REs) for safe sequestration within lipid droplets (LDs); and (2) the membrane-protective role of ACSL3, which incorporates monounsaturated fatty acids (MUFAs) into phospholipids (MUFA-PLs) to physically obstruct lipid peroxidation. Proteins involved in iron homeostasis (TFRC, SLC11A2, FTH1, FTL, HMOX1) and the antioxidant response (SLC7A11, GPX4, etc.) are indicated as key nodes regulating the sensitivity of this pathway. Red lines indicate inhibitory interactions, while black arrows denote enzymatic conversion or metabolic flux.

The lipidomic data associated with recovery identified genotype-associated differences in glycerophospholipid and neutral lipid profiles following atRAL exposure. Relative to WT cells after recovery, RDH12-expressing cells showed increased abundance of several PC and putatively annotated ether-linked PC species (**Figure 5G**). Additionally, PCs and DGs were differentially abundant (**Figure 5H**), including PC 34:1 and PC 36:7, whereas DG 30:3, DG 32:5, and DG 34:2 were less abundant (**Figure 5H**). In addition, we putatively annotated oxidized PC species in the recovery dataset based on the LIPID MAPS database. PC(18:0/Aze) was detected from the MRM transition *m/z* 694.5→184 in recovery samples but not in the acute dataset, whereas PC(16:0/8:(COOH)) with MRM transition *m/z* 652.5→184 was detected in both datasets but was not significantly altered between the conditions (**Figure 5H**). These observations show that oxidized-PCs were detectable after atRAL removal. The presence of late-stage lipid oxidation products indicates that RDH12-expressing cells were exposed to substantial levels of reactive oxygen species. While one would predict these should be higher in cells that do not express RDH12, the observation in RDH12 cells could simply reflect that these cells tolerated the stress and survived.

We also observed additional lipid class-specific differences between the recovery conditions. In WT and RDH12-expressing cells, recovery from atRAL was associated with a decreased abundance of PI and PE species relative to vehicle-treated cells (**Figure 5G**). Qualitative DBE analysis further indicated dose- and genotype-dependent differences in the DBE composition of the detected lipid precursors (**Figure S8A, B**). After recovery from 200 µM atRAL, RDH12-expressing cells showed a higher proportion of MUFA-containing DGs, whereas WT cells showed a higher proportion of PUFA-containing DGs. Nonetheless, PUFA-containing lipid species remained abundant in both recovery lipidomes, indicating that oxidizable lipids were still present after atRAL removal and are consistent with PUFA-containing lipids as critical components of cell membranes.

Together, the recovery proteomic and lipidomic datasets indicated that RDH12-expressing and WT cells occupy distinct molecular states after high-dose atRAL exposure. RDH12-expressing cells showed a stronger representation of proteins associated with microtubule and actin-cytoskeletal organization, and ferroptosis related signatures upon 200 µM atRAL, together with a lipid profile characterized by a higher abundance of selected PC and ether-linked PC species and a lower abundance of DG species relative to WT cells (**Figure 5G**, **Figure S8C-E**). In contrast, WT cells showed elevated proteins related to extracellular remodeling, cellular metabolism, and heme biosynthesis. These findings from the proteome and lipidome support the conclusion that RDH12 expression is associated with a distinct post-atRAL recovery state and indicate that RDH12 expression promotes organized membrane repair and structural recovery after atRAL-induced toxicity, whereas WT cells remain in a more disrupted stress response state. The emergence of ferroptosis-related signatures in the recovery dataset, together with oxidized phospholipid species detected in the lipidomic data, prompted us to further examine whether ferroptosis-associated mechanisms contribute to atRAL-induced toxicity.

### atRAL-induced Toxicity leads to ferroptosis-related cell death

Because we observed several oxidation-state- and metal-handling-related proteins and oxidized phospholipids in both the acute and recovery phases, we further examined the multiomics data with a focus on a ferroptosis-associated cellular response to atRAL toxicity. Ferroptosis is an iron-dependent form of regulated cell death driven by accumulation of reactive oxygen species and production of lipid peroxides (91). This process is promoted by the convergence of three features: disruption of the system Xc^−^/glutathione/GPX4 antioxidant axis, increased Fe2^+^ availability, and enrichment of polyunsaturated fatty acid-containing phospholipids (92). Ferritinophagy, a selective autophagic pathway mediated in part by NCOA4-dependent ferritin turnover, can further contribute to ferroptosis by mobilizing ferritin-stored iron and increasing the labile iron pool (93).

The proteomic response to atRAL exposure was consistent with the induction of oxidative and electrophilic stress processes (**Figure 6A**). Several proteins associated with KEAP1-NRF2-regulated stress adaptation were upregulated following atRAL exposure, including HMOX1, SLC7A11, GCLC, MGST2, and MGST3. SLC7A11 promotes cystine uptake, and GCLC catalyzes the rate-limiting step in glutathione synthesis (**Figure 6C**) (*58, 94*). Together, their increased abundance is consistent with a compensatory response to oxidative stress and an increased demand for glutathione-dependent antioxidant capacity. HMOX1, MGST2, and MGST3 further support the activation of heme- and electrophile-responsive detoxification pathways (*95, 96*).

In parallel, several antioxidant and peroxide-detoxifying proteins were downregulated, including GPX1, GPX4, TXNRD1, TXNRD2, GSS, and GSR. GPX4 is a central suppressor of ferroptosis in its use of glutathione to reduce phospholipid hydroperoxides. The decrease in GPX4, GSS, and GSR suggests that although cells increase upstream cystine import and partially activate glutathione synthesis, the downstream GPX/GSH-dependent lipid peroxide detoxification may be compromised during acute atRAL stress. This pattern is consistent with a compensatory response that may be insufficient to fully maintain lipid peroxide detoxification. In the recovery phase NRF2 targets were again elevated: HMOX1 remained higher in RDH12-expressing cells than in WT cells after 200 µM atRAL, and SRXN1 was detected only in RDH12-expressing cells. This time dependence suggests that the initial atRAL shock forces a transient collapse of cellular homeostasis, which the cell actively attempts to resolve through an NRF2-mediated electrophile detoxification program.

Furthermore, proteins involved in iron transport were upregulated, including TFRC, which mediates endocytosis of the transferrin-Fe^3+^-bound complex. SLC11A2 was upregulated and transports Fe^2+^ into the cytosol, contributing to the intracellular labile iron pool. SLC39A14 was also upregulated and transports non-transferrin-bound iron; it is known to be upregulated under pathological conditions (97). These changes are consistent with an increase in a free, redox-active Fe^2+^ pool that enables Fenton chemistry, leading to reactive oxygen species production and lipid peroxidation. Together, the data support altered iron homeostasis and ferroptosis-associated cell death mechanisms as a component of the atRAL response (**Figure 6C**). To determine whether ferroptosis contributes to atRAL-induced cell death and whether this effect varies by genotype, we compared the atRAL IC_50_ in wild-type (WT) and RDH12-expressing cells with and without 30 µM Ferrostatin-1 (Fer-1) pretreatment (**Figure 6B**). In WT cells, Fer-1 did not offer protection from atRAL: their IC_50_ was 133.7 µM (95% CI 127.0-140.4) without and 123.5 µM (95% CI 113.6-133.5) with the inhibitor, a 0.92-fold change indistinguishable from no effect (133 to 120 µM; Tukey HSD adj. p = 0.53). In contrast, Fer-1 pretreatment in RDH12-expressing cells increased IC_50_ from 220 µM (95% CI 213.4-226.9) to 352 µM (95% CI 330.8-372.2), a 1.60-fold increase (Tukey HSD, adj. p = 1.7 x 10^-4^). The genotype-by-ferrostatin interaction was significant (F(1,8) = 46.3, p = 1.4 x 10^-4^), and the separation between genotypes widened from 1.65-fold to 2.85-fold (**Figure 6B**). Collectively, these findings demonstrate that radical trapping protects only with sufficient RDH12 activity, indicating that lipid peroxyl radical propagation becomes a rate-limiting contributor to death only once aldehyde clearance is intact. In WT cells, ferroptosis does not represent the principal mechanism of cell death; instead, direct electrophilic modification of proteins and aminophospholipids likely predominates. These results suggest that RDH12 shifts the dominant mode of atRAL-induced cell death rather than acting synergistically with ferroptosis inhibition.

## Discussion

Retinaldehyde clearance is usually framed as a single enzymatic step: an NADPH-dependent reductase converts a reactive aldehyde into an inert alcohol, and the cell is spared. The genetics of RDH12 argue that this step is important (LCA13). Genetics does not explain what happens downstream of the enzyme or why photoreceptors cannot compensate. Here, we used paired proteomic and lipidomic profiling of a defined RDH12-expressing cell system, sampled during acute atRAL exposure and again after removal of atRAL, to place RDH12 in the sequence of events following an atRAL load. Three outcomes are observed. First, RDH12 acts as an upstream metabolic filter that raises the cytotoxic threshold for atRAL, rather than as a downstream survival factor. Second, acute atRAL exposure splits the glutathione axis, inducing the supply arm while depleting the terminal peroxidase arm and simultaneously expanding iron import, a combination that biases the cell toward lipid peroxidation. Third, RDH12’s role is most evident not during the insult but after it, when RDH12-expressing cells transition to a membrane- and cytoskeleton-repair state, whereas control cells remain in a remodeling and growth-factor-signaling state.

ER-localized RDH12 nearly tripled retinol output from an acute atRAL challenge while suppressing retinoic acid formation by more than 87% and shifting the 5 h IC_50_ from 133.7 µM to 220.2 µM. RDH12 does more than remove atRAL: it redirects retinaldehyde away from an irreversible oxidative branch. Without sufficient reductase capacity, endogenous aldehyde dehydrogenases oxidize atRAL, and the resulting retinoic acid is neither recyclable in the visual cycle nor inert. Therefore, reductive clearance is the only route that returns carbon to the retinoid pool, and RDH12 wins this competition by virtue of both catalytic efficiency and position, sitting on the ER membrane where atRAL partitions. The rightward shift in the dose–response curve is a threshold effect, not an on/off switch: RDH12-expressing cells still die; they simply require more aldehyde to do so. This is the expected signature of a rate-limiting detoxification enzyme and is consistent with what is observed in LCA3 patients with partially active RDH12 mutations.

The acute proteome shows that the cell reallocates rather than uniformly fails. Proteins increased at both doses were dominated by ER protein processing — the N-glycosylation and oligosaccharyltransferase machinery (ALG1, ALG11, DPAGT1, STT3A, STT3B, RPN1, RPN2, UGGT1) — together with solute carriers and glycerophospholipid enzymes, while proteins enriched in vehicle-treated cells were dominated by translation, transcription, and nucleic acid metabolism. This asymmetry, with 1,831 proteins decreased at 100 µM against 534 increased, reads as a translational and biosynthetic contraction with a selectively sustained secretory and membrane arm, the pattern expected of an integrated stress response. Notably, this occurred without the induction of canonical unfolded protein response chaperones. One interpretation is that the ER burden imposed by atRAL is not principally misfolded luminal proteins but membrane and glycosylation stress, to which the cell responds by reinforcing quality control at the point of entry rather than mounting a full UPR. The divergence within the cytoskeletal folding pathway, with the qualification that is dose-limited: CCT/TRiC subunits and tubulins rise at 100 µM and return to baseline at 200 µM, whereas prefoldin subunits fall at both doses. Folding capacity for existing clients is transiently reinforced while delivery of newly synthesized actin and tubulin, which depends on prefoldin, is curtailed along with translation.

The redox proteome contains the clearest mechanistic signal in the dataset: decoupling. The cystine–glutathione supply arm was induced — SLC7A11, GCLC, HMOX1, MGST2, and MGST3 all increased, the expected KEAP1–NRF2 output for an aldehyde — while the terminal peroxide-detoxifying arm fell, with GPX1, GPX4, GSS, GSR, TXNRD1, and TXNRD2 all decreased. Increased cystine import without commensurate GPX4 activity means the cell is buying a substrate it cannot fully use to scavenge phospholipid hydroperoxides. GPX4 is a specific enzyme that reduces peroxidized membrane lipids, and its loss triggers ferroptosis. Its decline in the presence of an active NRF2 program suggests that atRAL directly compromises this node. The second convergent input is iron. TFRC, SLC11A2, and SLC39A14 were all increased, spanning transferrin-bound uptake, endosomal Fe^2+^ export to the cytosol, and non-transferrin-bound iron import. The expansion of the labile iron pool alongside a weakened GPX4 arm is the canonical two-hit configuration for Fenton-driven lipid radical propagation, which arises from a single upstream insult. This provides a cell-autonomous mechanism for the recent demonstration that defective atRAL clearance drives photoreceptor degeneration through ferroptosis in mice (10), and it sits comfortably alongside reports that JNK inhibition attenuates atRAL-induced photoreceptor ferroptosis (19).

The lipidome after 200 µM atRAL treatment showed changes most directly traceable to atRAL chemistry. Acute exposure raised LPC 16:0, LPC 18:1, and PC 32:0, while select PE species were depleted. The PE decrease is notable because atRAL condenses with the primary amine of phosphatidylethanolamine to form N-retinylidene-PE, the precursor of retinal bisretinoids, so depletion of the PE pool is consistent with direct electrophilic consumption (98). Lysophospholipid accumulation is a general marker of phospholipid turnover and membrane stress.

The proteome showed a coincident increase in enzymes of associated with glyercophospholipid synthesis and remodeling, though the strength of evidence differs among them because the acute vehicle group was the least deeply sampled group in the dataset. LPCAT1, a reacylase favoring saturated and monounsaturated acyl-CoA donors, increased at both doses; the cytosolic calcium-dependent phospholipase A2 PLA2G4A increased at 200 µM, as did the lysosomal phospholipase A2 PLA2G15, indicating that the deacylation arm of the Lands cycle was engaged; and PTDSS1, which supports phosphatidylserine synthesis by base exchange, increased at both doses. A second and larger set of enzymes also scored as increased, comprising GPAT4 and AGPAT3/4/5 for phosphatidic acid, CHKA and ETNK1 in the Kennedy pathway, and DGKE linking diacylglycerol back to phosphatidic acid. Each of these was undetected in all three vehicle replicates, so should be read as directional rather than quantitative. Two enzymes central to the interpretation could not be evaluated at all: LPCAT3, which incorporates arachidonate and promotes ferroptosis through the ACSL4 axis, and the calcium-independent phospholipase A2 PLA2G6 were not detected in any sample. ACSL4 itself was quantified throughout and changed by less than 1.3-fold at both doses (99). Read together, the coincidence of elevated LPC with elevated LPCAT1 is consistent with reacylation skewed toward less oxidizable acyl chains, and the concurrent rise in PC 32:0 would be compatible with LPCAT1-mediated reacylation of LPC 16:0 with palmitoyl-CoA, though our MRM-based annotation does not allow us to demonstrate that route definitively (100). PNPLA6 was also scored as increased at 200 µM, but on a vehicle baseline quantified in only one of three replicates, and neither dose reached significance. In retinal pigment epithelial cells PNPLA6 acts as a phospholipase B that deacylates PC and then hydrolyzes the resulting LPC to glycerophosphocholine, which re-enters PC synthesis through the Kennedy pathway (70). If the increase is genuine, it would favor enhanced engagement of this PC regeneration loop over net LPC production. These data are consistent with an attempted membrane-remodeling response, but they do not establish that repair outpaces damage.

Acyl composition analysis was dose-dependent and did not resolve into a single pattern. DG precursors showed a higher proportion of PUFA-containing species at 100 µM as did TG precursors at 200 µM, while ceramide precursors shifted toward saturated species and PC composition remained comparatively stable. Diversion of polyunsaturated acyl chains into neutral lipid pools is a described means of limiting the peroxidizable membrane fraction, and the stability of PC composition does not argue against it. The PC pool is substantially larger than the DG and TG pools, so a proportionally large enrichment in the latter would produce only a small compositional shift in the former. However, these data cannot distinguish diversion from de novo synthesis or from uptake of exogenous polyunsaturated fatty acids. As the MRM transitions used here targeted unoxidized precursors, PUFA species consumed by peroxidation would be lost from the panel rather than be observed as a compositional change. Therefore, neutral-lipid PUFA enrichment is compatible with, but not definitive evidence for, protective sequestration of oxidizable acyl chains (101). Separately, four ceramides and dihydroceramide species increased at 100 µM but not at 200 µM. Sphingolipids participate in stress signaling, autophagy, and ferroptosis-associated pathways, so this may represent one component of the lipid stress response (76, 102).

The recovery experiment is where the two genotypes separate most informatively, and the separation is qualitative rather than one of degree. After recovery from 200 µM atRAL, RDH12-expressing cells were enriched for microtubule and actin cytoskeletal organization and for an NRF2-target module (HMOX1, SRXN1) (84) and carried higher levels of PC and putatively ether-linked PC species with lower DG. In contrast, control cells accumulated CCN1, CCN2, FN1, WNT5A, ERBB2, FGFR2, SPRY2, and ATF3, which are matricellular, ERK/MAPK, and injury-response proteins, along with a heme biosynthetic module (ABCB10, UROS, IBA57). The RDH12 state reflects a cell restoring structure and membrane composition after a survivable insult, while the control state reflects a cell still signaling injury. Plasmalogens and other ether phospholipids act as sacrificial antioxidants and markers of peroxisomal lipid synthesis; however, ether-lipid biosynthesis has been reported to sensitize cells to ferroptosis (103, 104). Therefore, their elevation in the protected genotype cannot be assigned a direction of effect based on abundance alone. The same caution applies to oxidized species: the exclusive detection of PC(18:0/Aze) in recovery samples indicates that lipid oxidation products persist after aldehyde removal. However, as noted, the cells in which they are detected are, by definition, survivors, and their appearance in RDH12-expressing cells reflects sampling from a living population rather than a greater oxidative burden.

The ferrostatin-1 experiment identifies RDH12 within this sequence. Pre-treatment with 30 µM ferrostatin-1 for 24 h left the atRAL IC_50_ of control cells essentially unchanged (133.7 µM to ∼120 µM), while raising that of RDH12-expressing cells from 220.2 µM to ∼352 µM, widening the genotype separation from 1.65-fold to 2.85-fold. Therefore, radical trapping is protective only in the context of the RDH12 background. Because ferrostatin-1 acts solely by intercepting lipid peroxyl radicals, its failure to protect control cells indicates that the killing these cells experience is not primarily driven by that chemistry. The most probable interpretation is that the two interventions act on different segments of one pathway, and that the segment blocked by ferrostatin-1 becomes rate-limiting only once the upstream segment is relieved. In cells without adequate reductase capacity, atRAL persists at concentrations at which cell death is dominated by direct electrophilic chemistry, that is, Schiff base and Michael adduction of proteins and aminophospholipids, and the consequent failure of the thiol- and selenol-dependent enzymes documented above, which a lipid radical scavenger cannot intercept (105). Once RDH12 lowers the aldehyde burden below this threshold, iron-dependent lipid peroxide propagation carries out residual killing, and radical trapping becomes effective. Consistent with this interpretation, a recent independent study demonstrated that atRAL exposure induces the canonical ferroptosis signature, including iron accumulation, lipid peroxidation, and SLC7A11/GPX4 suppression, in retinal pigment epithelial cells and in a mouse model of impaired atRAL clearance. In both systems, Fer-1 rescued cell viability (106). Thus, ferroptosis is not the universal executioner of retinaldehyde toxicity, but the executioner that emerges once aldehyde clearance is intact, reframing the relationship between the two and consistent with the observation of ferroptotic degeneration in models that retain partial clearance capacity (10, 19). The clinical corollary is testable: ferroptosis-directed therapy would be predicted to benefit patients retaining residual RDH12 activity and to underperform in those with complete loss unless combined with restored clearance. It also cautions against transferring ferroptosis rescue results between models with different aldehyde-handling capacities.

However, several limitations bound these conclusions. HEK293T cells provide a tractable and genetically clean background in which the presence or absence of a single reductase can be interrogated; however, they lack the defining features of a photoreceptor: an outer segment, DHA- and VLC-PUFA-rich disc membranes, phototransduction-scale metabolic demand, and the endogenous visual cycle that generates atRAL continuously in situ. Therefore, our system models the biochemistry of retinaldehyde detoxification, not the cell biology of the photoreceptor, and the acyl-chain conclusions should be revisited in a membrane environment with the relevant lipid composition. The experimental conditions of short exposure at high concentrations, in contrast to long exposures, should also be considered carefully.

Notwithstanding these caveats, the data support a coherent temporal model with therapeutic implications. Because RDH12 acts before the oxidative cascade begins, restoring reductase capacity is the intervention with the broadest reach, which underlies gene replacement approaches for RDH12-associated dystrophy. When clearance cannot be restored, the model proposes downstream nodes that are pharmacologically addressable and that our data show cells are engaged in radical trapping, iron chelation, GPX4 stabilization, and modulation of membrane acyl composition toward monounsaturated species. The persistence of the RDH12 advantage despite ferroptosis inhibition also suggests that these downstream strategies would be additive to, not redundant with, restoring the enzyme itself. Extending this framework to photoreceptor-organoid models expressing patient-derived RDH12 variants, with direct measurement of lipid peroxidation and labile iron, is the logical next step toward testing whether the sequence defined here operates in the retina.

## Acknowledgements

The authors thank the Purdue Proteomics Facility and Metabolite Profiling Facility and its staff, in particular, Dr. Christina Ferreira for data collection and consultation in interpretation.

## Lead Contact

Bryon S. Drown

## Data Availability

Proteomics data has been deposited to PRIDE under the ProteomeXchange accession **PXD080689** (reviewer token: 7B6YZYEhu4GY). Lipidomics data has been deposited in the NIH Common Fund’s National Metabolomics Data Repository (NMDR) website, the Metabolomics Workbench, https://www.metabolomicsworkbench.org, where it has been assigned Study ID **ST005027**. The data can be accessed directly via its Project DOI: http://dx.doi.org/10.21228/M8F86Z

## Author Contributions

KV.D.: Conceptualization, Methodology, Validation, Formal analysis, Investigation, Resources, Writing – original draft. M. R. M.: Conceptualization, Methodology, Validation, Formal analysis, Investigation, Writing – original draft. R.V.: Investigation. D.A.: Formal analysis. B.S.D.: Conceptualization, Supervision, Formal analysis, Data curation, Funding acquisition, Writing – original draft, Writing - Review and Editing. R. S.: Conceptualization, Supervision, Funding acquisition, Writing - Review and Editing. O. B.: Methodology, Validation, Formal analysis, Investigation, Writing – Review and Editing. N.Y.K.: Funding acquisition, supervision, Writing - review and editing.

## Funding and Additional Information

This work was supported by an internal award at Purdue University as part of the Life Science Summit. The Bruker Tims-TOF^HT^ used to collect some of the data in this study was acquired by the Purdue Proteomics core facility via a National Institute of Health S10 (S10OD032364) award. N.Y.K. was supported by R01AR076924.

## Supplemental Information

This article contains supporting information:

• Figure S1. Proteomic characterization demonstrating the impact of RDH12 overexpression.

• Figure S2. Proteomic data coverage and quality.

• Figure S3. RDH12 abundance and relative rank across acute and recovery atRAL conditions.

• Figure S4. Missingness and imputation quality control for proteomic analysis.

• Figure S5. Gene ontology enrichment analysis for acute proteomics data.

• Figure S6. Lipid quality control analysis of datasets.

• Figure S7. Gene ontology enrichment of recovery proteome data.

• Figure S8. Recovery Lipidomics analysis.

• SI Table 1. DIA window scheme for proteomics data collection (CSV)

• SI Table 2. Protein group evidentiary support tiers (CSV)

• SI Table 3. Counts of peptides supporting each identified protein group in Exploris dataset (CSV)

• SI Table 4. Counts of peptides supporting each identified protein group in timsTOF dataset (CSV)

• SI Table 5. Protein abundance across acute and recovery datasets (CSV)

• SI Table 6. Protein differential abundance analysis in acute and recovery data sets (CSV)

• SI Table 7. Lipid abundance in acute dataset (CSV)

• SI Table 8. Lipid abundance in recovery dataset (CSV)

• SI Table 9. Lipid differential abundance analysis in acute dataset (CSV)

• SI Table 10. Lipid differential abundance analysis in recovery dataset (CSV)

• SI Data 1. Extracted ion chromatograms supporting identification of all protein groups with only one proteotypic peptide (PDF)

• SI Data 2. Extracted ion chromatograms supporting identification of protein groups named in main text with only one proteotypic peptide (PDF)

