## Supplemental Figures for "Multiomic Profiling Links RDH12-Dependent Retinaldehyde Detoxification to Membrane Remodeling and Ferroptosis": Supplemental_Figures.pdf

**Supporting Information**

**List of materials**

- **Figure S1.** Proteomic characterization demonstrating the impact of RDH12 overexpression.
- **Figure S2.** Proteomic data coverage and quality.
- **Figure S3.** RDH12 abundance and relative rank across acute and recovery atRAL conditions.
- **Figure S4.** Missingness and imputation quality control for proteomic analysis.
- **Figure S5.** Gene ontology enrichment analysis for acute proteomics data.
- **Figure S6.** Lipid quality control analysis of datasets.
- **Figure S7.** Gene ontology enrichment of recovery proteome data.
- **Figure S8.** Recovery Lipidomics analysis.
- **SI Table 1**. DIA window scheme for proteomics data collection (CSV)
- **SI Table 2**. Protein group evidentiary support tiers (CSV)
- **SI Table 3**. Counts of peptides supporting each identified protein group in Exploris dataset (CSV)
- **SI Table 4**. Counts of peptides supporting each identified protein group in timsTOF dataset (CSV)
- **SI Table 5**. Protein abundance across acute and recovery datasets (CSV)

- **SI Table 6**. Protein differential abundance analysis in acute and recovery data sets (CSV)
- **SI Table 7**. Lipid abundance in acute dataset (CSV)
- **SI Table 8**. Lipid abundance in recovery dataset (CSV)
- **SI Table 9**. Lipid differential abundance analysis in acute dataset (CSV)
- **SI Table 10**. Lipid differential abundance analysis in recovery dataset (CSV)
- **SI Data 1**. Extracted ion chromatograms supporting identification of all protein groups with only one proteotypic peptide (PDF)
- **SI Data 2**. Extracted ion chromatograms supporting identification of protein groups named in main text with only one proteotypic peptide (PDF)

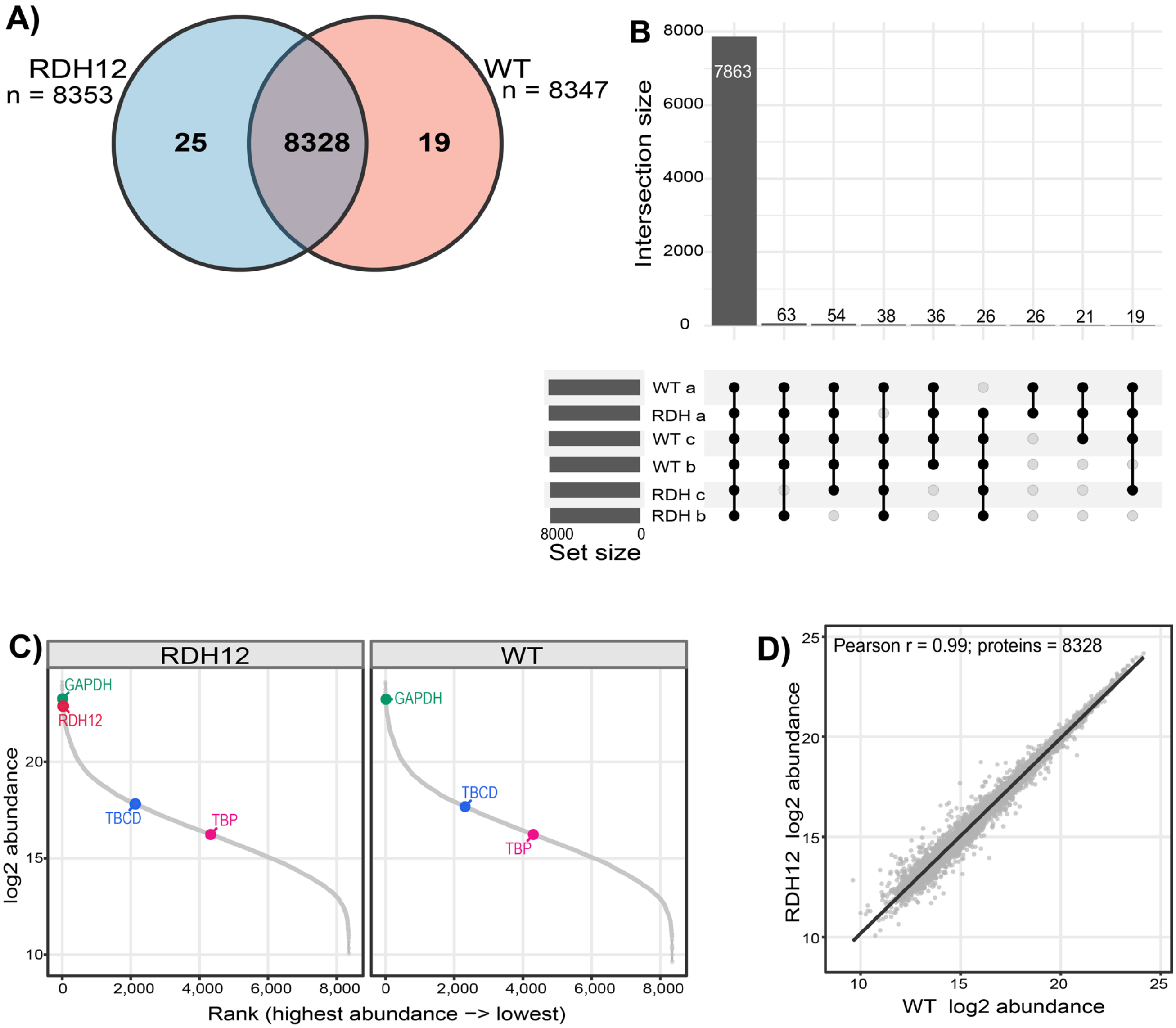


**Figure S1. Proteomic characterization demonstrating the impact of RDH12 overexpression.** (A) Venn diagram showing the overlap of protein groups identified in RDH12-expressing and WT HEK293T cells. Of the 8,372 proteins detected, 8,328 were shared between the two cell lines, with 25 proteins unique to RDH12-expressing cells and 19 unique to WT cells. (B) UpSet plot showing the overlap of protein identifications across biological replicates (a-c), demonstrating high reproducibility and extensive overlap of detected proteins. (C) Rank-abundance plots of protein groups detected in RDH12-expressing and WT cells. Proteins are ranked according to mean $\log_2$ abundance. Representative proteins across a range of abundances are highlighted, including GAPDH, TBCD, TBP, and RDH12. RDH12 was detected exclusively in the RDH12-expressing cell line. (D) Correlation of mean $\log_2$ protein abundances for the 8,328 protein groups shared between RDH12-expressing and WT cells with a Pearson correlation of r = 0.99.

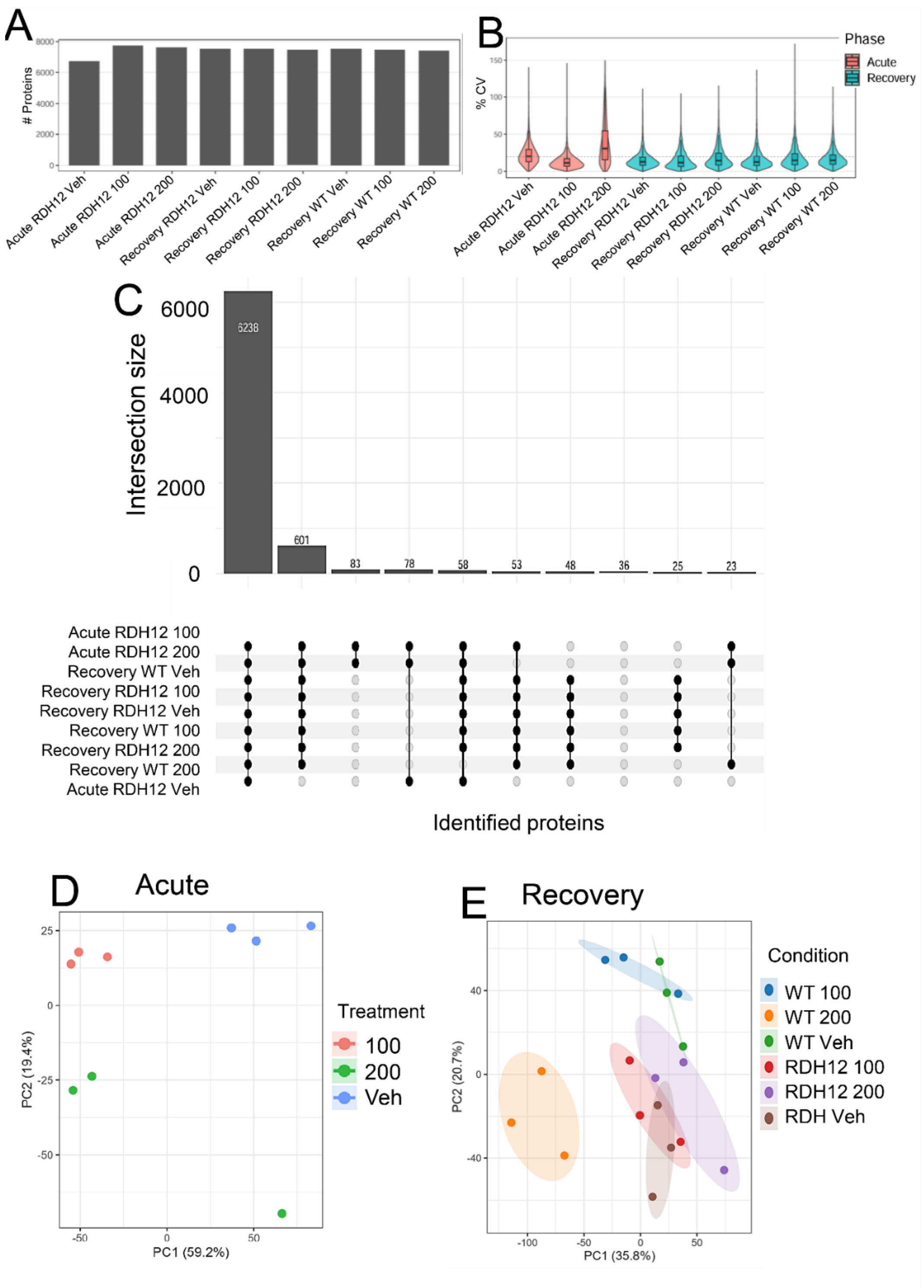


**Figure S2. Proteomic data coverage and quality.** (A) Number of proteins identified in each sample. (B) Distribution of protein-level coefficient of variation (CV) values across conditions, calculated from non-imputed LFQ intensities across biological replicates. Acute samples are shown in red, and recovery samples are shown in blue. (C) UpSet plot showing the intersection of the identified proteins across samples and experimental conditions (D) Principal component analysis (PCA) of the acute RDH12 proteomics dataset prior to imputation. (E) PCA of the recovery proteomics dataset prior to imputation.

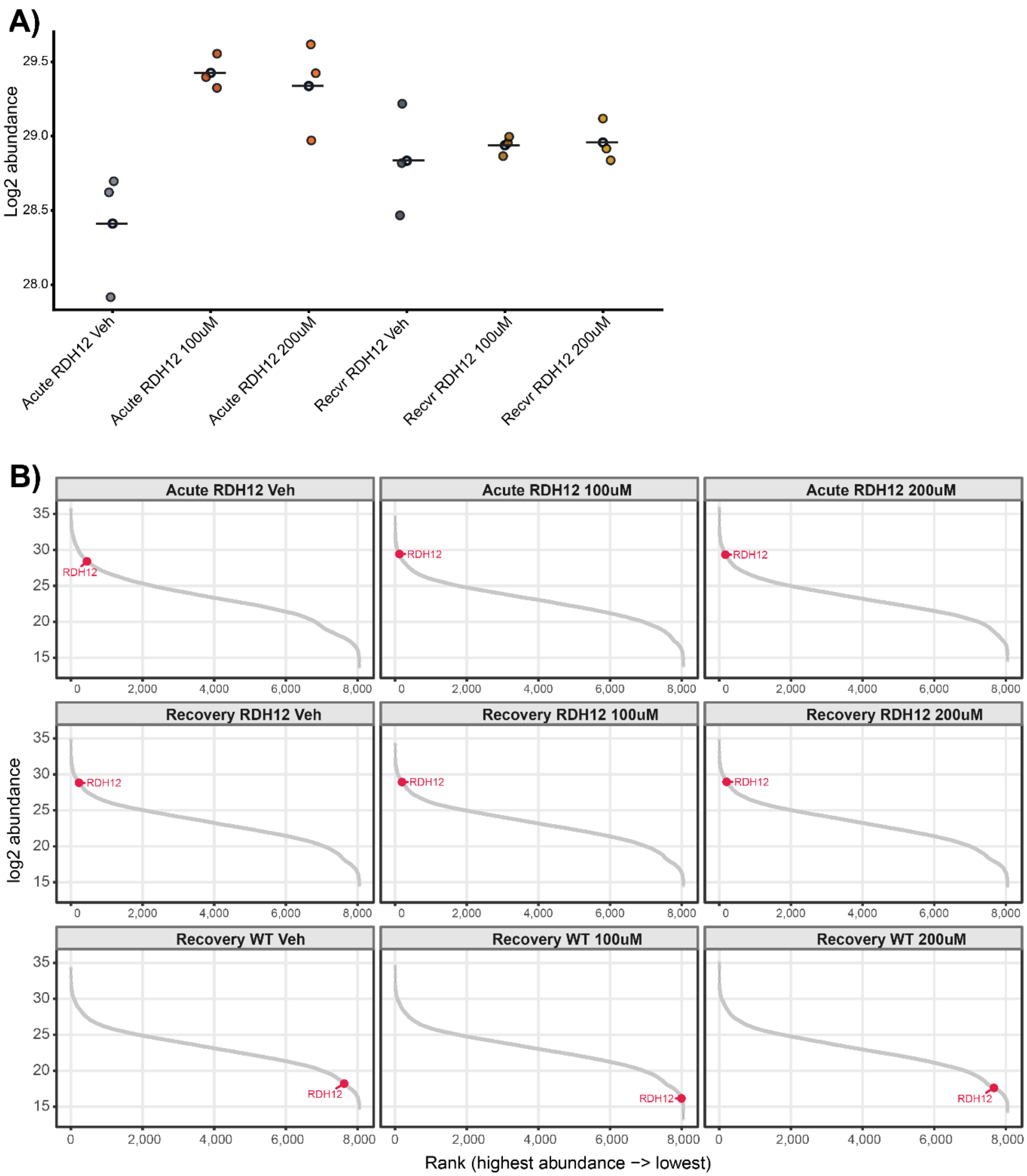


**Figure S3. RDH12 abundance and relative rank across acute and recovery atRAL conditions.** (A) Log2-transformed RDH12 protein abundance in RDH12-expressing HEK293T cells following acute 5 h exposure to vehicle, 100 μM, or 200 μM atRAL and after 24 h recovery following the same treatments. Points represent individual replicates, and horizontal lines indicate the mean abundance. (B) Rank-abundance distributions of detected proteins within each acute and recovery condition, ordered from highest to lowest abundance. RDH12 is highlighted in red to show its relative abundance rank within each sample. RDH12 was not detected in WT recovery samples, its displayed position in these panels was generated from imputed missing values and should not be interpreted as endogenous RDH12 expression in WT cells.

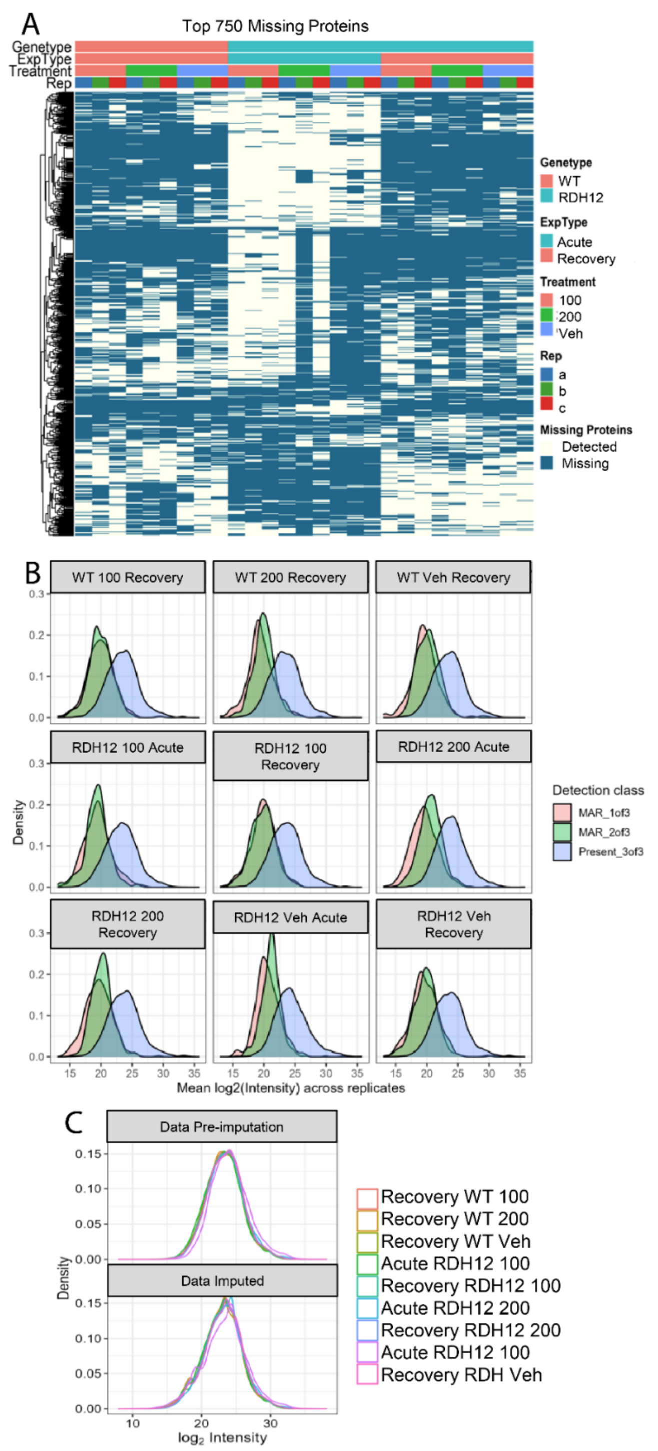


**Figure S4. Missingness and imputation quality control for proteomic analysis.** (A) Heatmap of missing protein values across samples for the top 750 proteins with the highest missingness. Columns are annotated by genotype, experiment type, treatment, and biological replicates. (B) Density distributions of the mean log2 protein intensity by missingness class within each condition. Proteins detected in all three biological replicates generally showed higher mean intensities than proteins missing in one (MAR 1 of 3) or two replicates (MAR 2 of 3), consistent with missingness being enriched among lower-abundance proteins. (C) Density distributions of log2 protein intensities before and after mixed imputation.

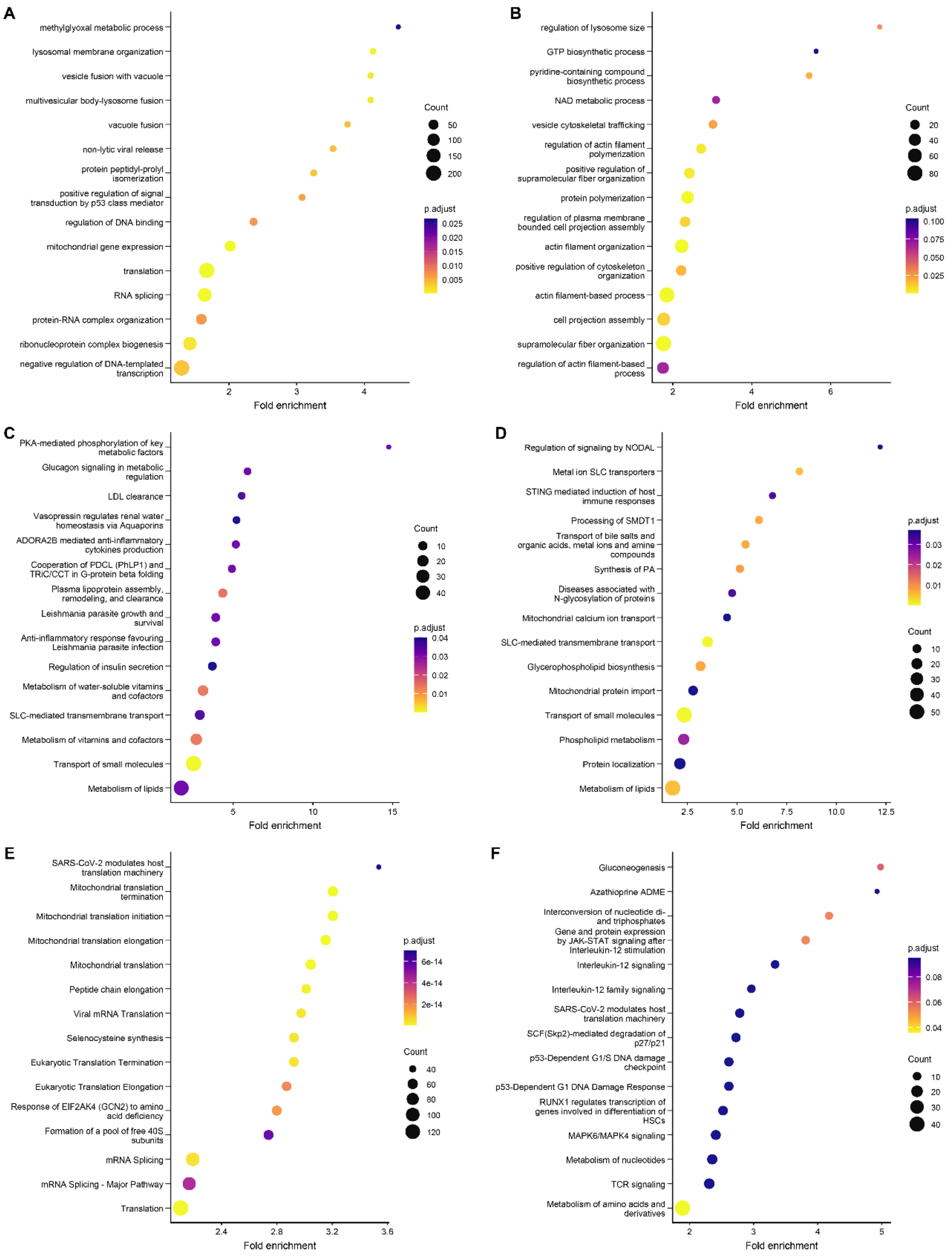


**Figure S5**. **Gene ontology and pathway enrichment analysis for acute proteomics data.** Biological process GO enriched terms from proteins more abundant in vehicle-treated cells relative to atRAL-treated cells at (A) 100 µM and (B) 200 µM. (C) Reactome pathway enrichment from proteins more abundant in 100 µM atRAL-treated cells relative to vehicle-treatment. (D) Reactome pathway enrichment from proteins more abundant in 200 µM atRAL-treated cells relative to vehicle-treatment. (E) Reactome pathway enrichment from proteins more abundant in vehicle cells relative to 100 µM atRAL-treated cells. (F) Reactome pathway enrichment from proteins more abundant in vehicle cells relative to 200 µM atRAL-treated cells.

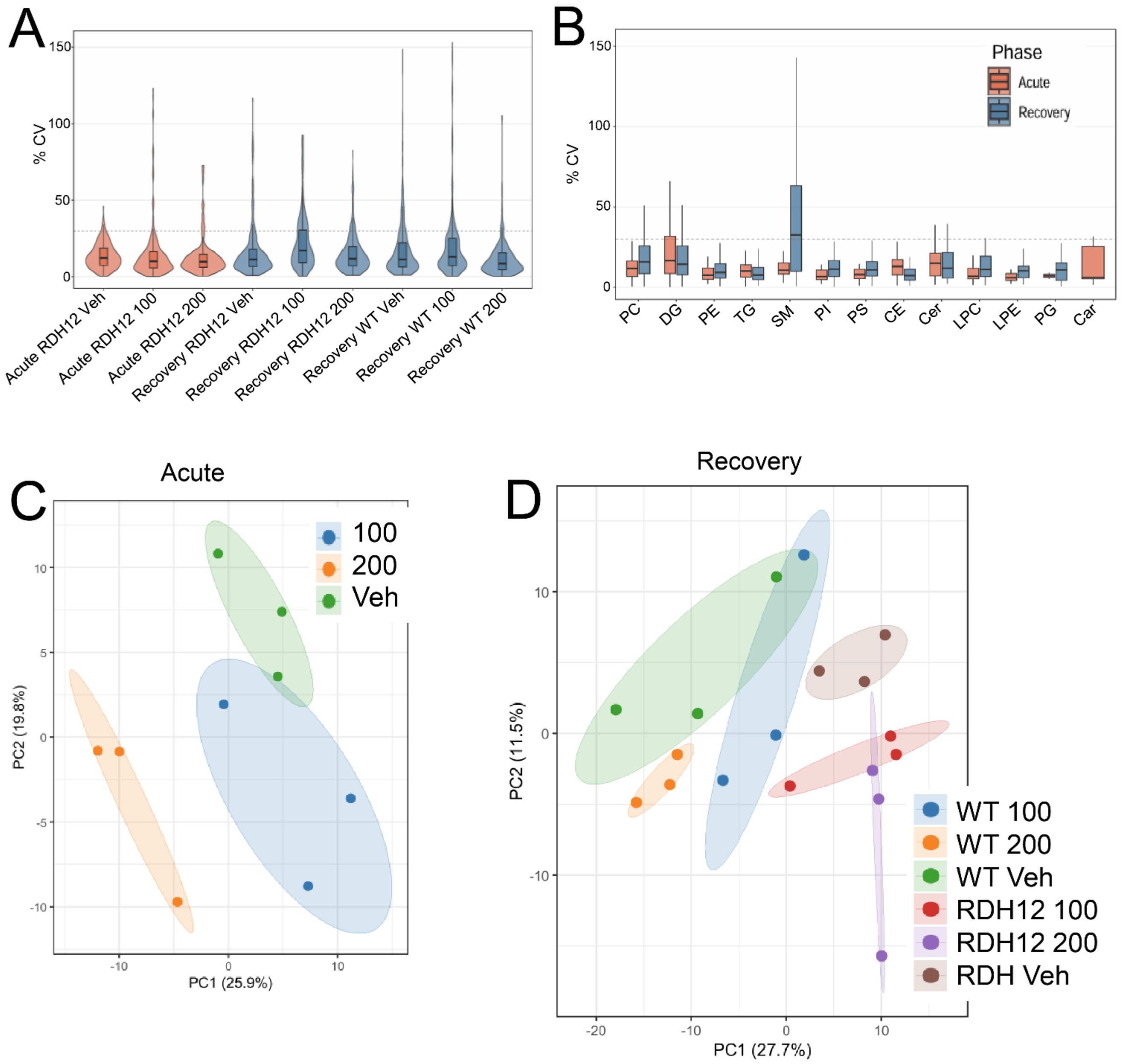


**Figure S6. Lipid quality control analysis of datasets**. (A) Distribution of lipid coefficient of variation (CV) values under acute and recovery conditions. The CVs were calculated from the raw lipid intensity values across biological replicates within each condition. The dashed line indicates a 30% CV. (B) Distribution of lipid CV values grouped by lipid class. The dashed line indicates a 30% CV. (C) Principal component analysis (PCA) of the acute RDH12 lipidomics dataset, showing separation among vehicle, 100 μM atRAL, and 200 μM atRAL conditions. (D) PCA of the recovery lipidomics dataset, showing separation by genotype and treatment condition across RDH12-expressing and WT cells.

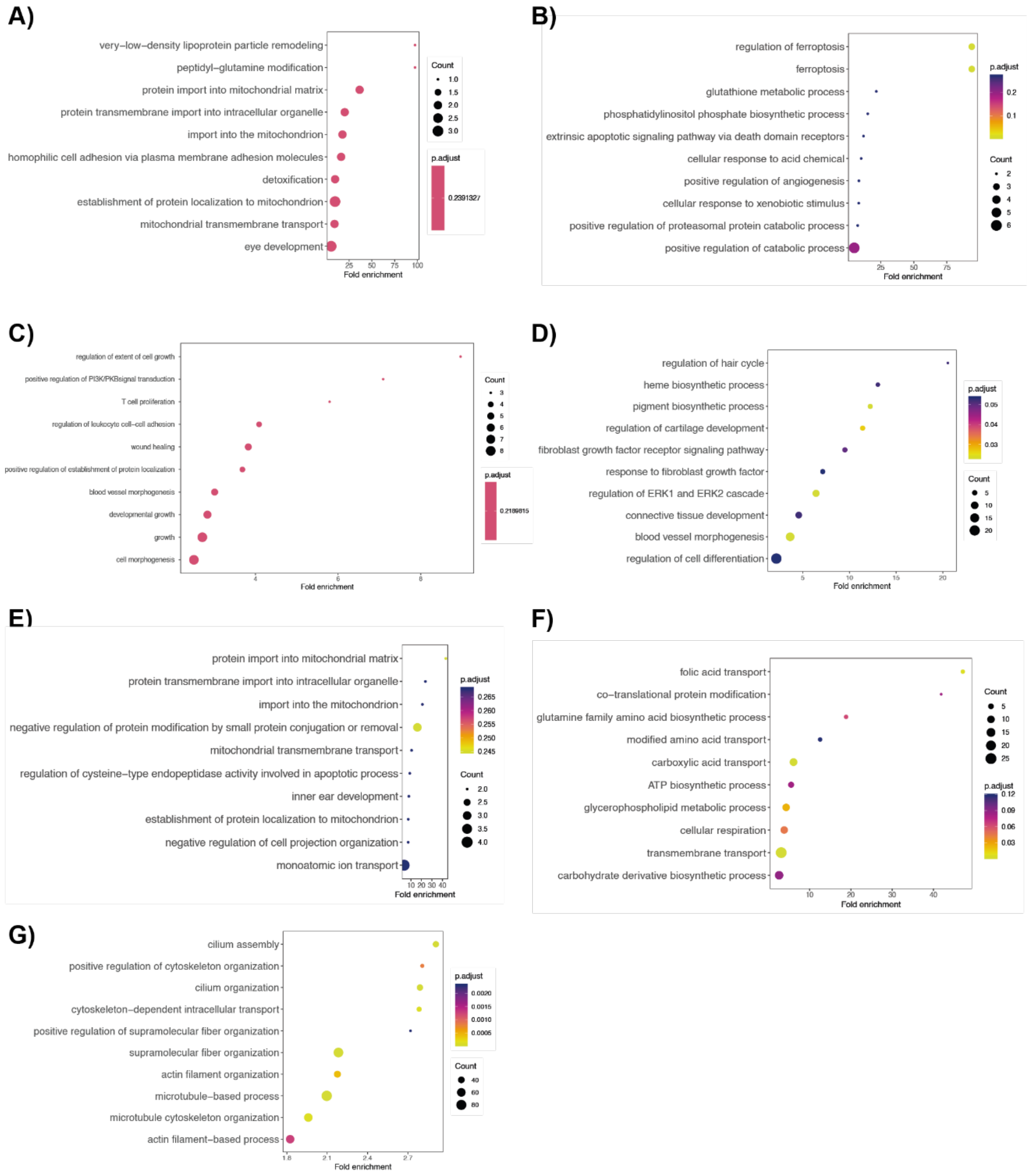


**Figure S7. Gene ontology enrichment of recovery proteome data.** (A) BP GO analysis from proteins more abundant in RDH12-expressing cells in Vehicle-treatment relative to 100 μM atRAL. BP GO analysis *of* (B) proteins more abundant in RDH12-expressing cells in 200 μM atRAL treatment relative to Vehicle, and (C) proteins more abundant in RDH12-expressing cells in Vehicle treatment relative to 200 μM atRAL treatment. (D) BP GO analysis of proteins more abundant in WT cells relative to RDH12-expressing cells after 200 μM atRAL. BP GO analysis of proteins more abundant in WT cells in 100 μM atRAL treatment relative to Vehicle. BP GO analysis from (F) shows proteins more abundant in WT cells in 200 μM atRAL-treatment relative to Veh, and (G) shows proteins more abundant in WT cells in Vehicle-treatment relative to 200 μM atRAL-treatment.

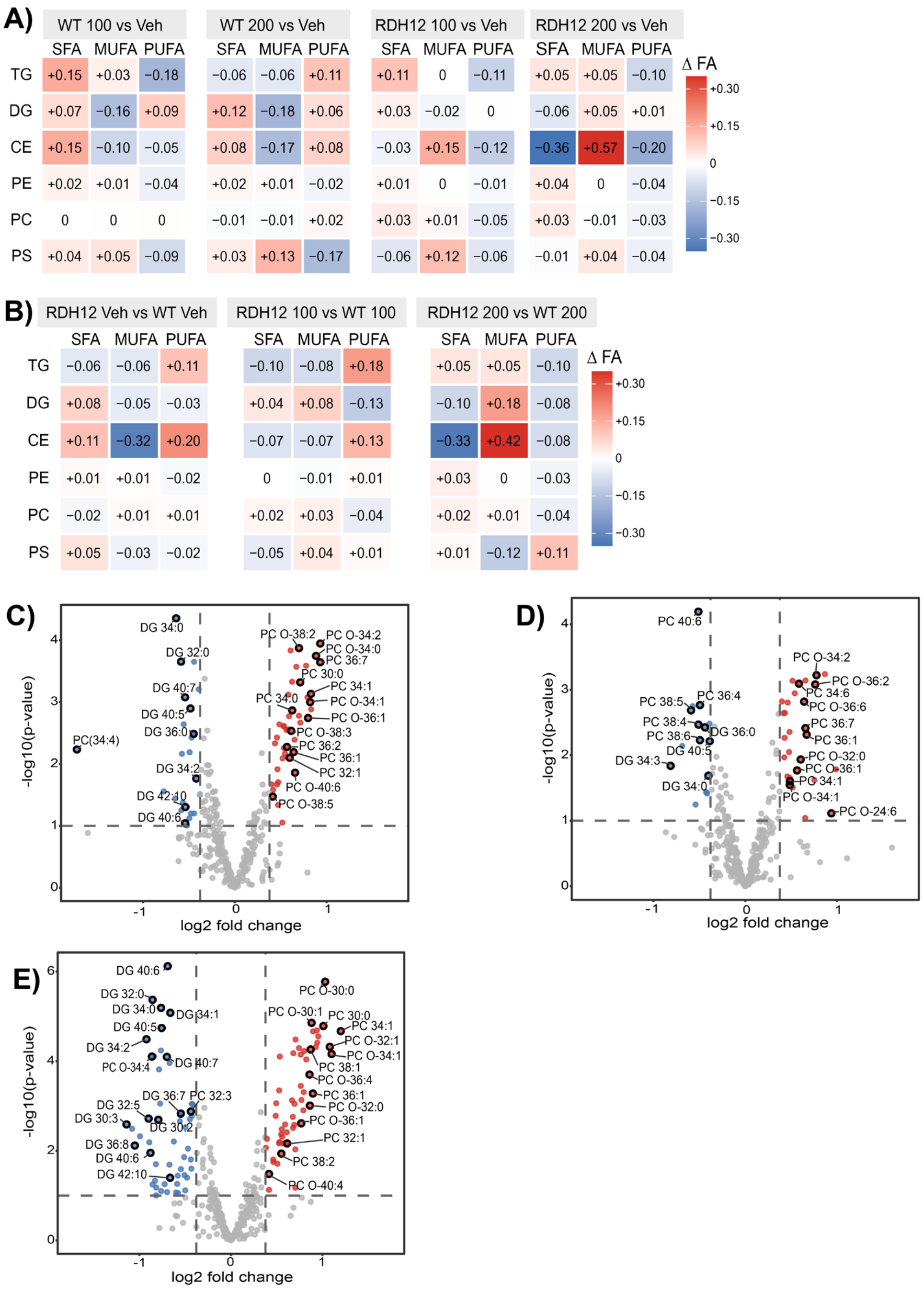


**Figure S8. Recovery Lipidomics analysis**. Qualitative DBE analysis further indicated dose- and cell-dependent differences in the composition of the detected lipid precursors. The heatmap shows the change in double bond equivalent (DBE) composition of distinct detected lipid precursors for (A) dose vs Veh comparisons in WT and RDH12-expressing cells and (B) for cell-type comparisons within each dose. Within each lipid class and treatment group, precursors were grouped as SFA (0 DBE), MUFA (1 DBE), or PUFA (≥2 DBE), and the proportions of identifications for each class were calculated. Tile values indicate the change in precursor proportion. Red indicates a higher proportion, and blue indicates a lower proportion within that comparison. Volcano plots comparing WT vs RDH12 at each dose (C) Vehicle, (D) 100 µM and (E) 200 µM. Vertical dashed lines indicate ±1.3-fold change, and the horizontal dashed line indicates a nominal p-value cutoff of 0.1
